# Chronological lifespan extension and nucleotide salvage inhibition in yeast by isonicotinamide supplementation

**DOI:** 10.1101/2021.07.11.451986

**Authors:** Agata I. Kalita, Christopher T. Letai, Elisa Enriquez-Hesles, Lindsey N. Power, Swarup Mishra, Shekhar Saha, Manikarna Dinda, Dezhen Wang, Pankaj K. Singh, Jeffrey S. Smith

## Abstract

Isonicotinamide (INAM) is an isomer of the NAD^+^ precursor nicotinamide (NAM) that stimulates the enzymatic activity of Sir2, an NAD^+^-dependent histone deacetylase from the budding yeast, *Saccharomyces cerevisiae*. Supplementing INAM into growth media promotes the replicative lifespan (RLS) of this single cell organism by maintaining intracellular NAD^+^ homeostasis. INAM also extends yeast chronological lifespan (CLS), but the underlying mechanisms remain largely uncharacterized. To identify interacting genes, a chemical genomics screen of the yeast knockout (YKO) collection was performed for mutants sensitized to growth inhibition by INAM. Significant Gene Ontology (GO) terms included transcription elongation factors, metabolic pathways converging on one-carbon metabolism, and de novo purine biosynthesis, collectively suggesting that INAM may perturb nucleotide metabolism. Indeed, INAM caused dose-dependent depletion of intracellular cytidine, uridine and guanosine, ribonucleosides derived from the breakdown of nucleotide monophosphates by a set of nucleotidases (Phm8, Sdt1, Isn1) or the alkaline phosphatase Pho8. Direct inhibition of recombinant Sdt1 and Phm8 nucleotidase activity by INAM was confirmed *in vitro*, as was inhibition of alkaline phosphatase activity. Each of these enzymes can also convert nicotinamide mononucleotide (NMN) to nicotinamide riboside (NR), consistent with an accumulation of NMN and NAD^+^ upon inhibition by INAM. Taken together, the findings suggest a model whereby partial impairment of nucleotide salvage pathways can trigger a hormetic stress response that supports enhanced quiescence during chronological aging.

## Introduction

Aging is a multifaceted process leading to decreased function over time, and ultimately, death of the cell or organism. Despite extraordinary variance in organism lifespans, many of the proteins and pathways involved in aging are evolutionarily conserved (1, 2), making research in short-lived, experimentally feasible model organisms applicable to human aging. For example, budding yeast (*Saccharomyces cerevisiae*) has played a major role in the identification and characterization of several key longevity or health span factors, including the NAD^+^-dependent histone deacetylase Sir2 (reviewed in (3)), the founding member of the conserved sirtuin protein family (4). Sir2 is critical for sustaining the replicative potential of yeast mother cells, also known as their replicative lifespan (RLS), primarily through stabilizing the repetitive rDNA locus (5). In contrast, shown in further work in yeast, Sir2 limits the long-term survival of non-proliferating cells in stationary phase, also known as chronological lifespan (CLS), due to specific Lys-514 deacetylation (inactivation) of the rate-limiting gluconeogenesis protein Pck1 (6). Gluconeogenesis improves ethanol and acetate utilization and elevates glycogen/trehalose storage, thus enhancing stress resistance and CLS (7).

Sirtuin-mediated lysine deacetylation produces nicotinamide (NAM) as a byproduct, which can feed-back and inhibit the reaction if not recycled by the NAD^+^ salvage pathway (8-10). NAM supplementation at high concentrations therefore phenocopies the effect of deleting the *SIR2* gene, reducing RLS and extending CLS (9, 11). Biochemically, NAM inhibits deacetylation activity by regenerating acetyllysine and NAD^+^ through a base exchange reaction with the peptidyl-imidate intermediate (12). Isonicotinamide (INAM), a non-reactive isostere of NAM (Fig. 1A), can relieve the Sir2 NAM inhibition and stimulate deacetylation activity by blocking the NAM binding pocket (13). Supplementing INAM into media therefore enhances heterochromatic gene silencing and RLS in a Sir2-dependent manner (13, 14). However, rather than shortening CLS, as would be predicted for enhanced Sir2 activity, INAM extends CLS independently of Sir2 (15), suggesting the involvement of potentially novel CLS regulatory mechanisms.

**Fig. 1.**
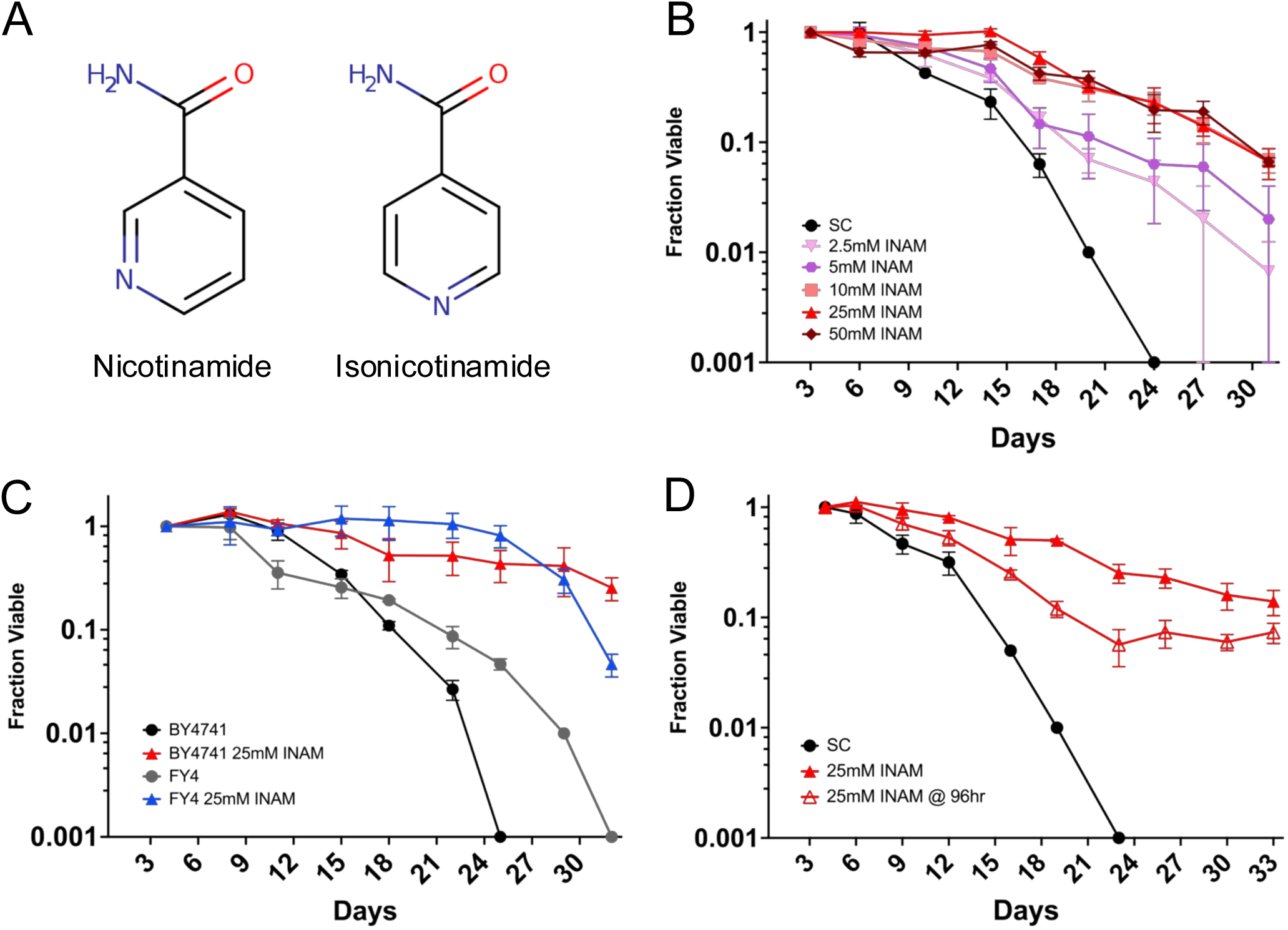
INAM extends chronological lifespan. **A)** Chemical structures of nicotinamide and isonicotinamide. **B)** CLS assays with increasing dosage of INAM added to BY4741 at the time of inoculation. **C)** CLS assay comparing the effect of 25 mM INAM on the auxotrophic strain BY4741 and prototrophic strain FY4, added at inoculation. **D)** CLS of BY4741 when 25 mM INAM is added 96 hr after inoculation. Error bars indicate standard deviations (n=3 biological replicates). Oasis 2 statistics for CLS assays are provided in Table S5.

While INAM stimulates the HDAC activity of recombinant Sir2 in vitro (13), it also promotes Sir2-dependent transcriptional silencing and RLS in vivo by maintaining intracellular NAD^+^ homeostasis when NAD^+^ precursors (NAM or nicotinic acid) are limiting (14). INAM is not a direct NAD^+^ precursor, so its effects on NAD^+^ homeostasis likely involve the modulation of one or more steps of the de novo synthesis or salvage pathways. Consistent with this idea, NAM supplementation during chronological aging maintains the expression of several genes in these pathways when cells enter stationary phase, including the nicotinic acid/nicotinamide mononucleotide adenylyltransferase *NMA1* (14). INAM also upregulates the expression of genes involved in cell wall biosynthesis and remodelling, likely related to concurrent protection from acetic acid toxicity (15). However, the biochemical targets underlying these longevity-associated phenotypes remain unknown. Identifying such targets could provide important CLS mechanistic insights.

To identify possible INAM-targeted cellular processes, we utilized an unbiased chemical synthetic lethality screening approach with the yeast knockout (YKO) strain collection. We hypothesized that concentrations of INAM that extend CLS may moderately inhibit cellular processes would become growth inhibitory when combined with deletion of a gene from an interacting pathway or process. Deletion mutants that were growth-inhibited by sub-lethal INAM concentrations clustered into several functional Gene Ontology (GO) terms, including transcription elongation factors. This was similar to the synthetic lethal pattern observed with mycophenolic acid (MPA), a compound that inhibits IMP dehydrogenase and depletes the guanine nucleotide pool (16, 17). However, rather than specific reduction of guanine nucleotide biosynthesis, we found that INAM acutely impaired the salvage of ribonucleoside monophosphates CMP, GMP, UMP, and the NAD^+^ precursor, nicotinamide mononucleotide (NMN) to their respective nucleosides. Ultimately, this limited the overall nucleotide triphosphate pool as cells entered stationary phase. We propose a model whereby moderately limiting the nucleotide pool through inhibition of nucleotidase activity by INAM induces a beneficial hormesis response that improves CLS.

## Results

### INAM-induced CLS extension is titratable and independent of auxotrophies

To begin characterizing the mechanism of INAM-induced CLS extension, we tested for dosage effects on the YKO parental strain BY4741. Previous in vivo experiments with INAM were performed at 25 mM, a concentration that promotes Sir2-dependent transcriptional silencing (13). In Fig. 1B, concentrations ranging from 2.5 to 50 mM were supplemented into CLS cultures at the time of inoculation. Lifespan plateaued at 10 to 25 mM, while 50 mM still extended CLS despite causing reduced viability at day 3. To rule out contributions of the auxotrophic mutations in BY4741 to CLS extension, we showed that 25 mM INAM extended CLS of the prototrophic progenitor strain FY4 ((18); Fig. 1C). Additionally, 25 mM extended CLS of BY4741 when added to cultures at day 4, after the cells entered stationary phase (Fig. 1D). The effect was not as strong as supplementing at the time of inoculation but still suggested the underlying mechanisms impact both proliferating and quiescent cells.

### INAM extends CLS independently of sirtuins

We previously found that INAM supplementation prevented NAD^+^ depletion that normally occurs as yeast cells enter stationary phase (14, 19). Despite elevated NAD^+^ being associated with increased *SIR2* function, the associated CLS extension did not require *SIR2* (15). Significant functional redundancy exists between Sir2 and its paralog Hst1 (20-22), so to rule out contributions from Hst1 or the other Sir2 homologs, we tested whether INAM extended CLS of a mutant lacking all 5 sirtuins, *SIR2*, *HST1*, *HST2*, *HST3*, and *HST4* (4). CLS of the quintuple mutant was reduced compared to the control, but 10 mM INAM still significantly extended it (Fig. 2A), suggesting that while improved NAD^+^ homeostasis may play a role in the CLS benefit, sirtuins are not involved.

**Fig. 2.**
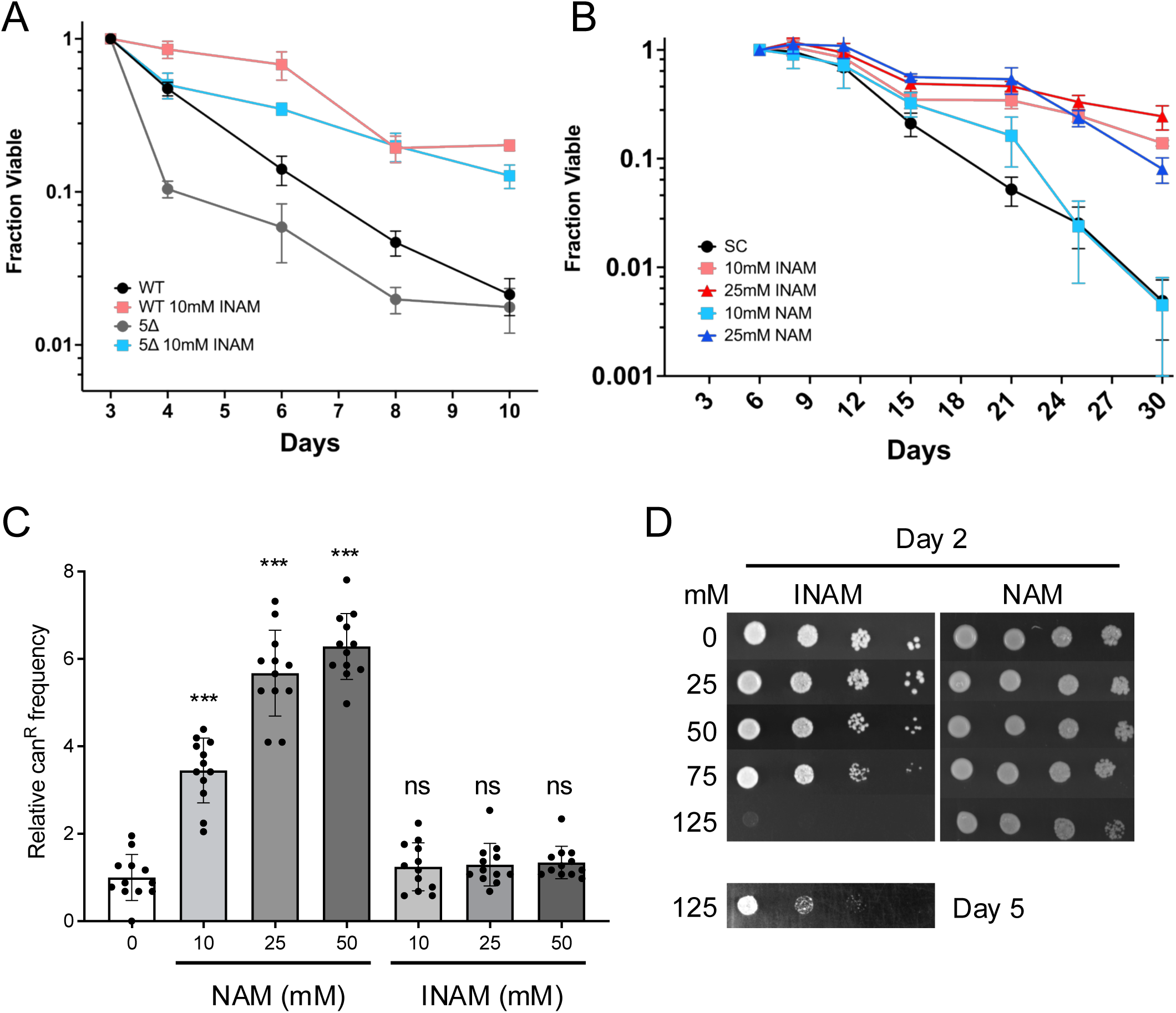
INAM effects on CLS are independent of sirtuins. **A)** CLS assay demonstrating that 10mM INAM extends CLS of a quintuple sirtuin (*sir2Δ, hst1Δ, hst2Δ, hst3Δ, hst4Δ*) mutant. B) CLS assay comparing the effects of INAM and NAM supplemented to BY4741 at 10 or 25mM. Oasis 2 statistics for CLS assays are provided in Table S5. **C)** Relative mutation frequency caused by overnight growth on NAM or INAM at indicated concentrations. The average frequency for the control without supplements was set at 1.0. Error bars indicate standard deviation (n=12), and asterisks indicate p-values <0.001 when compared to the control (1-way Anova test). **D)** Growth of BY4741 with increasing concentrations of INAM or NAM on SC media. 10-fold serial dilutions of cells. The 125 mM INAM plate was also incubated to day 5.

The lack of sirtuin involvement suggested that despite being a sirtuin inhibitor, NAM may also extend CLS due to its structural similarity with INAM. We previously reported that 5 mM NAM did not extend CLS when added to cultures at the time of inoculation (23), but it was subsequently shown to extend CLS when added during the diauxic shift (11). NAM at 5 mM inhibits Sir2, Hst1, Hst2, and Hst4, while 25 mM NAM is required to inhibit Hst3 and ensure the elimination of all sirtuin function (9, 24). In Fig. 2B, we observed that 25 mM NAM significantly extended CLS of BY4741, but 10 mM NAM had little effect compared to 10 mM INAM, indicating that INAM is the more potent isomer by this measure. NAM also has mutagenic potential at these concentrations due to the inhibition of Hst3 and Hst4, which function as H3K56 deacetylases in the maintenance of genome stability (25, 26). Indeed, NAM supplementation significantly increased mutation frequency of the endogenous *CAN1* reporter gene (canavanine resistant colonies) as previously reported (26), while INAM at the same concentrations had no effect (Fig. 2C). INAM is therefore a non-mutagenic CLS booster in comparison to NAM. At concentrations above 50 mM, however, INAM inhibited cell growth while NAM did not (Fig. 2D). This led to the hypothesis that lifespan extension at lower INAM concentrations could be due to mild perturbation of one or more activities/pathways that function in cell proliferation.

### A chemical genomic screen for INAM-sensitive deletion strains

To identify genes that buffer against INAM-impaired cellular processes, the *MAT*a haploid YKO strain collection was arrayed onto rectangular OmniTray YPD agar plates, and the colonies were then pinned onto synthetic complete (SC) agar plates containing 0, 25, 50, 75, or 125 mM INAM (Fig. S1A). After 2 days incubation (3 days for 125 mM), the plates were imaged and growth of each mutant colony on the INAM-containing plates was quantitated relative to SC control plates using SGAtools (27). An example of the growth inhibition by INAM is shown for *tho2Δ* in Fig. S1B (center colony, yellow arrow). Growth of all strains was strongly reduced at 125 mM. The full screen was independently performed twice, showing moderate but significant correlation in SGAtools-generated fitness scores, as shown for 75 mM INAM in Fig. S1D (R = 0.42; p<0.00001). We conservatively focused on mutants with a fitness score lower than -0.3 (difference visible by eye; (27)) in both replicates (Tables S1 and S2, Fig. S1C inset). Fewer mutants met this stringent criterion at 50 mM (173), and 25 mM (150), which was expected as the degree of pathway disruption should be weaker (Fig. S1D). Only 39 mutants were identified from all four concentrations, though there were greater overlaps for any two concentrations (Fig. S1D). Approximately 50% of the 326 mutants chosen as sensitive to 125 mM INAM at day 3 did not overlap with mutants from the other concentrations (Fig. S1D), suggesting a high number of false positives for this highest concentration. We therefore focused Gene Ontology (GO) enrichment analysis on a collection of 341 genes identified from at least one of the 25, 50-, or 75-mM conditions (Table S1). Several ‘Biological Process’ GO terms with significant p-values were related to transcriptional regulation, autophagy and vacuolar transport, inositol phosphate biosynthesis, and chromatin remodelling (Table S3). ‘Cellular Component’ GO terms covered the INO80 and SWR1 chromatin remodelling complexes, transcriptional elongation factors, endosomes and the ESCRT complex (Table S3). INAM sensitivity was confirmed for 57 of 61 re-tested deletion mutants (Fig. S2 and S3), indicating the cut-off score of -0.3 was highly stringent. Based on predictions from the enriched GO terms and literature searches, 22 additional mutants related to the GO terms were directly tested and confirmed as INAM-sensitive (Fig. S4).

### Disruption of serine, glycine, threonine, or cysteine metabolism confers sensitivity to INAM

Intriguing metabolism-related GO terms enriched in the 25, 50, or 75 mM INAM-sensitive mutant lists were *de novo* IMP biosynthesis (*ADE1*, *ADE4*, *ADE6*, *ADE5,7*), threonine metabolic process (*HOM2*, *HOM3*, *THR4*, and *GLY1*), and serine family amino acid biosynthetic process (*CYS4*, *GLY1*, *HOM2*, *HOM3*, *SER1*, *SER2*) (Table S3). Fig. 3A depicts connections between these enzymes, with the points of convergence at serine and glycine, amino acids that fuel one-carbon (1C) metabolism for utilization in purine and thymidylate (dTMP) synthesis, methionine/SAM regeneration, amino acid synthesis, and redox maintenance (28). Serine metabolism has been linked to CLS regulation with serine supplementation extending CLS (29, 30), and *SER1* identified as a major QTL for genetic CLS variation among strain backgrounds (31). The serine/glycine/threonine pathways were also reported as a metabolic axis for longevity regulation based on multi-omics analyses in mice (32). While INAM clearly interacts with this metabolic axis, deletion mutants for the core one-carbon metabolism genes were not isolated from the sensitivity screen, with the exception of *ade3Δ*, which is also critical for the de novo purine synthesis pathway.

**Fig. 3.**
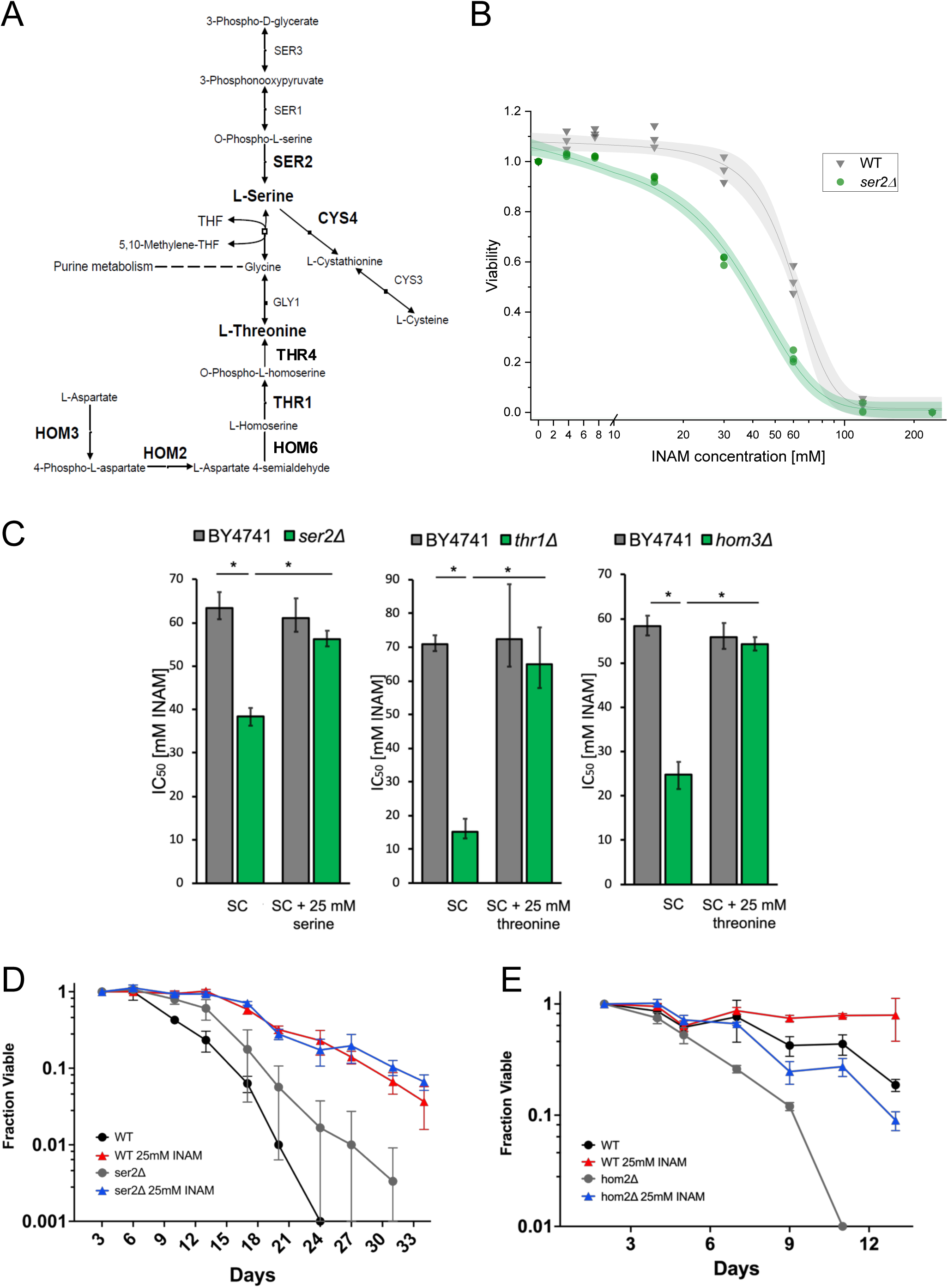
Serine and threonine biosynthesis mutants are sensitive to INAM. **A)** A diagram illustrating the synthesis pathways of serine, aspartate, threonine, and cysteine, as well as the integration into one-carbon metabolism and purine metabolism. Deleted genes conferring INAM sensitivity are in bold. **B)** Representative quantitative dose-response curve illustrating the relative viability of BY4741 (WT) and the *ser2Δ* mutant at a range of INAM concentrations. Dots and inverted triangles represent values obtained from three biological replicates with sigmoid curves fitted to the data using a dose response function and the Levenberg Marquardt iteration algorithm. Shaded areas around the curve are 95% confidence bands. **C)** Mean INAM IC_50_ values for BY4741 (WT), *ser2Δ, thr1Δ, and hom3Δ* strains calculated from the equations from the fitted curves as in panel B. Error bars represent 95% confidence intervals. **D)** CLS assay showing moderate lifespan extension of *ser2Δ* mutant, and full extension by 25 mM INAM. **E)** CLS assay showing shortened lifespan of a *hom2Δ* mutant that is partially restored by 25 mM INAM. Oasis 2 statistics for CLS assays are provided in Table S5.

Numerous metabolic enzymes ‘moonlight’ with biochemical functions separate from their annotated catalytic activity (33). We therefore wanted to confirm that the serine or threonine metabolism mutants were INAM-sensitive due to a serine or threonine deficit, respectively. To quantify the effects on growth, IC_50_ values for INAM were calculated from small liquid cultures in 96-well plates, as shown for *ser2Δ* in Fig. 3B. Supplementing 25 mM serine indeed restored normal growth to the *ser2Δ* mutant and, similarly, 25 mM threonine restored growth to the more severe *thr1Δ* and *hom3Δ* mutants (Fig. 3C). Even though these mutants were sensitive to high concentrations of INAM, their CLS was still extended by 25 mM INAM (Fig. 3D and E; data shown for *ser2Δ* and *hom2Δ*). Note that the *hom2Δ* mutant was short-lived compared to WT, indicating a role for this pathway in supporting CLS. Poor growth of the *thr1Δ* mutant was incompatible with CLS assays. Taken together, these results suggest that INAM does not function by directly modifying serine, threonine, or glycine metabolic pathways. Rather, defects in availability of serine, glycine, or threonine via their biosynthesis or import impair growth when INAM perturbs a functionally linked cellular process.

### Mutants in SWR1 complex subunits and other transcriptional elongation factors are sensitive to INAM

Prominent non-metabolic GO terms included those related to transcriptional regulation and chromatin remodeling, especially at the level of transcriptional elongation (Table S3). Among these were multiple subunits of the SWR1 complex, a conserved ATP-dependent chromatin remodeler that exchanges H2A core histones in nucleosome octamers for the H2A.Z variant encoded by *HTZ1* (Fig. 4A; (34, 35)). Exchange of H2A with H2A.Z (Htz1) by SWR1 impacts multiple steps of gene expression, including facilitating progression of polymerase through nucleosomes during transcription elongation (36). Mutants for 5 SWR1 subunits (*bdf1Δ, vps72Δ, swc5Δ, swr1Δ, yaf9Δ*) were identified from the screen at 25, 50, or 75 mM. Deletions for three additional annotated SWR1 mutants (*arp6Δ*, *swc3Δ*, *swc7Δ*) were tested separately. The *swc7Δ* mutant was the only SWR1 candidate we tested that was not INAM-sensitive (Fig. 4A), most likely because the Swc7 subunit is not required for H2A.Z binding or histone replacement activity (37). The *htz1Δ* strain was also isolated from the screen and confirmed as sensitive at 25 mM (Fig. 4B). Several other factors associated with elongation were identified from the INAM sensitivity screen, including subunits of the THO complex, the elongator complex, and the C-terminal domain kinase complex (CTDK-I), suggesting that INAM impairs one or more cellular processes that support transcriptional elongation.

**Fig. 4.**
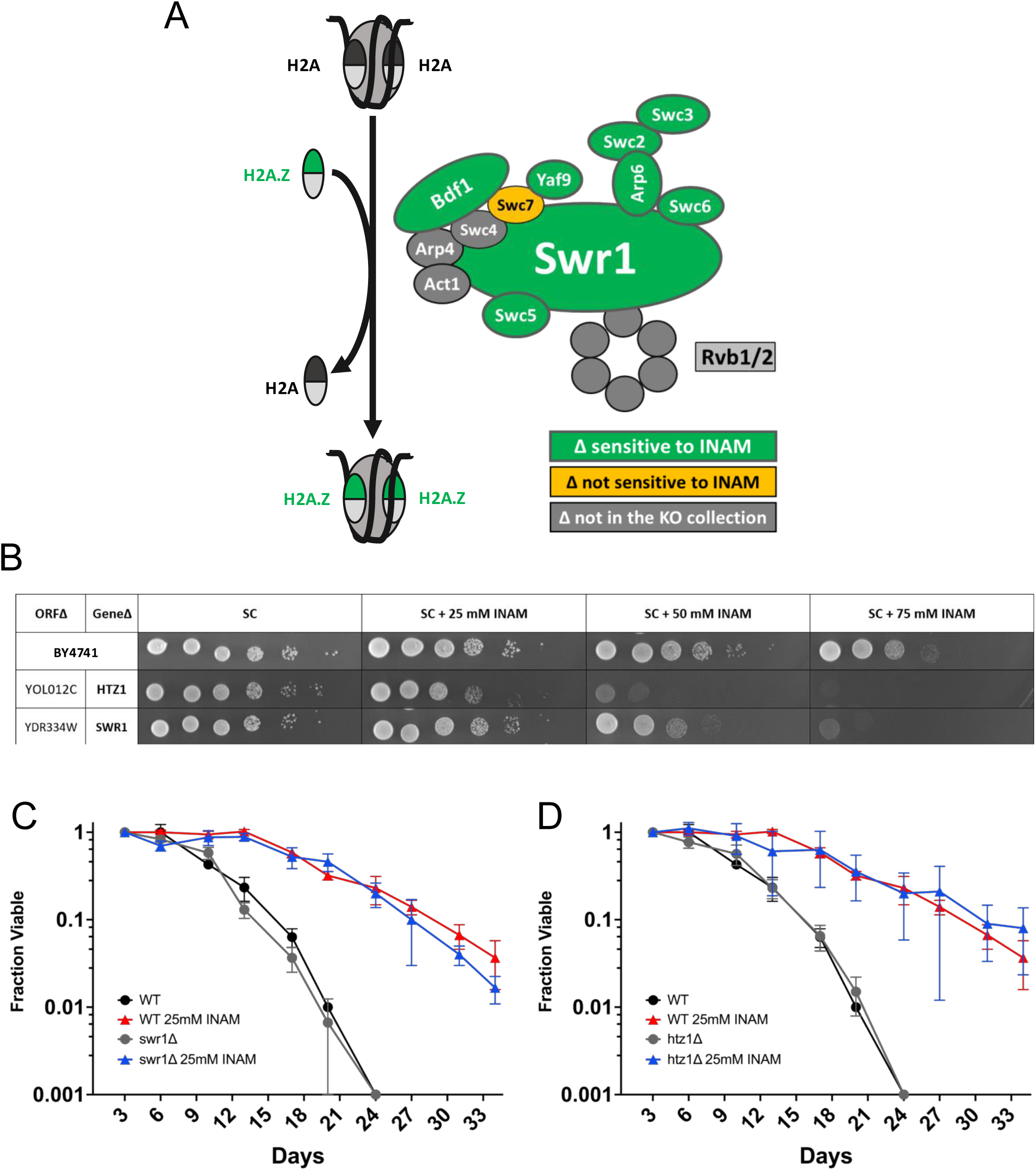
Strains defective for the H2A.Z deposition by SWR1 are sensitive to INAM. **A)** Illustration depicting the SWR1 activity of H2A/H2A.Z (Htz1) exchange, as well as subunit composition and SWR1 complex. Green indicates subunit deletions that were sensitive to INAM and yellow indicates not sensitive. **B)** Spot test growth assay showing confirmation of INAM sensitivity of *htz1Δ* and *swr1Δ* strains from the YKO collection. The WT strain is BY4741. **C)** CLS assay of BY4741 and *swr1Δ* mutant strains supplemented with 25 mM INAM. **D)** CLS assay of BY4741 and *htz1Δ* strains supplemented with 25 mM INAM. Controls were supplemented with equal volume of sterile water instead of INAM stock solution. Oasis 2 statistics for CLS assays are provided in Table S5.

SWR1 subunit mutations were previously shown to extend CLS of a prototrophic strain in nitrogen-rich media and to prevent the CLS extension induced by nitrogen-poor media, a form of caloric restriction (38). We therefore tested if deleting *SWR1* or *HTZ1* would prevent CLS extension induced by INAM. Surprisingly, neither mutant affected CLS with or without 25 mM INAM supplementation (Fig. 4C and D), suggesting any transcriptional or chromatin defects associated with SWR1 were not significantly contributing to longevity under the conditions of this study. Instead, synthetic lethality of INAM with elongation factor mutants, as well as de novo IMP biosynthesis pathway (*ade*) mutants, pointed toward possible alterations in nucleotide metabolism upon INAM supplementation that could impact CLS.

### INAM acts synergistically with mycophenolic acid (MPA)

Yeast mutants defective for transcriptional elongation are sensitive to mycophenolic acid (MPA) and 6-azauracil (6AU) (39). MPA is a non-competitive inhibitor of inosine 5’-monophosphate (IMP) dehydrogenase (IMPDH) (16), the enzyme that converts IMP to XMP during *de novo* synthesis of guanine nucleotides (Fig. 5A). 6AU is converted into 6-azaUMP, which also inhibits IMPDH (Imd2), as well as orotic acid decarboxylase (Ura3), the last step of UMP biosynthesis (40). The resulting guanine ribonucleotide pool depletion, and UTP depletion in the case of 6AU, impairs transcription elongation and inhibits growth when combined with mutations that slow elongation (41, 42). Comparing the INAM-sensitive mutants with previously identified MPA-sensitive mutants (39), there was significant overlap between the datasets, with 45.1% (46/102) of MPA-sensitive mutants also identified as INAM-sensitive (Fig. 5B). Among the overlaps were *htz1Δ* and multiple transcriptional elongation factors, suggesting that INAM is modifying ribonucleotide pools like MPA. Dose response growth curves of BY4741 with combinations of INAM and MPA in liquid cultures revealed strong synergistic growth inhibition (peak ZIP score of 9.86) at relatively low concentrations of each compound that had no effects individually (Fig. 5C), implying the two compounds were likely impacting growth through different mechanisms.

**Fig. 5.**
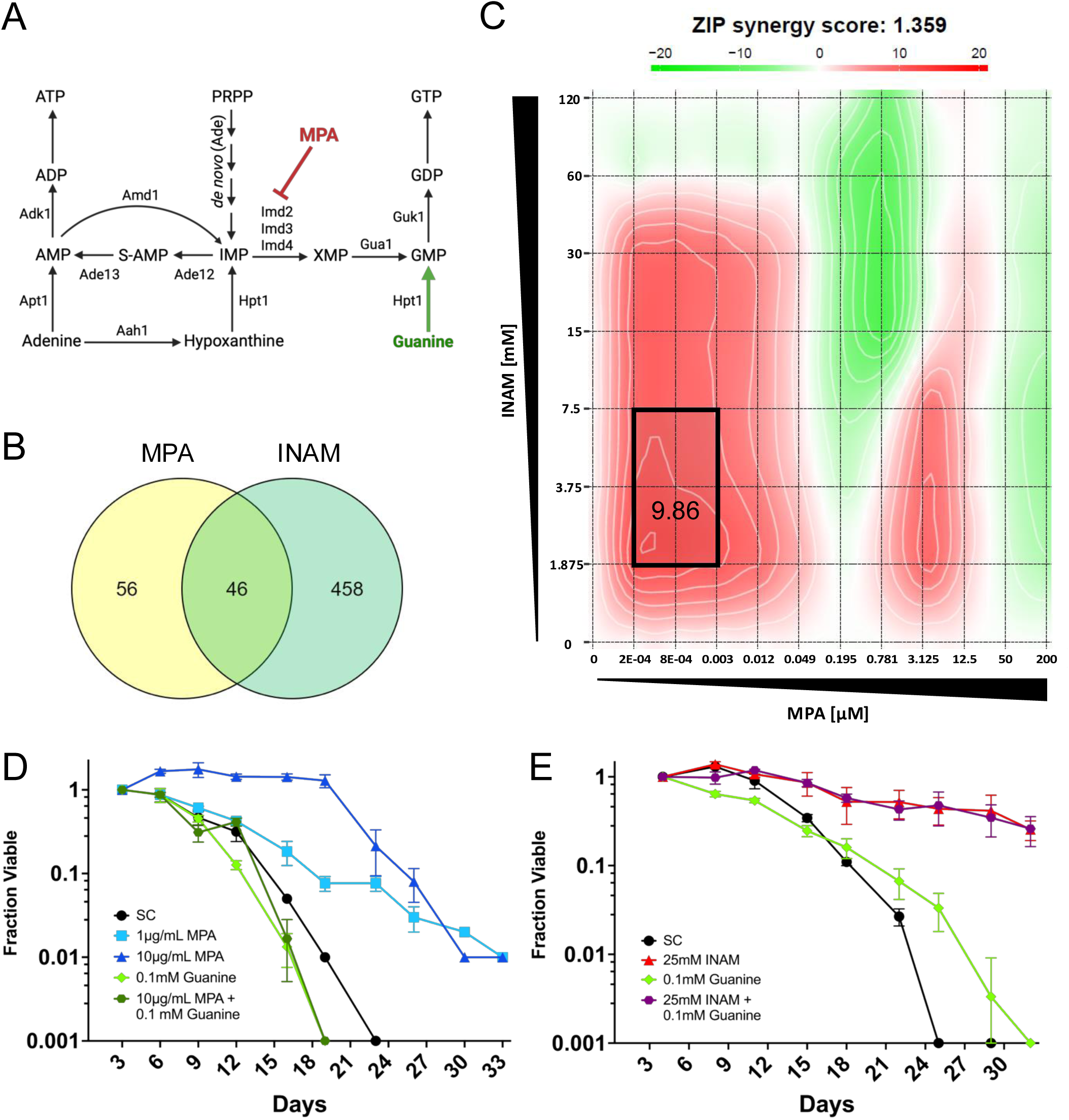
Mycophenolic acid and INAM operate through related but distinct mechanisms. **A)** Illustration of MPA inhibiting IMP dehydrogenase (Imd2) to reduce guanylic nucleotide synthesis. Guanine salvage via Hpt1 (green) compensates to support GMP, GDP, and GTP homeostasis. **B)** Venn diagram depicting a partial overlap between INAM-and MPA-sensitive deletion strains identified in a chemogenomic screen and confirmed by spot assays. **C)** Synergy map depicting interaction between INAM and MPA in reducing cell growth. Graph is representative of three independent experiments. **D)** CLS assay with BY4741 showing CLS extension by MPA supplementation (1 or 10 µg/ml), and suppression of the effect by 0.1 mM guanine. **E)** CLS assay with BY4741 showing INAM extension of CLS is that is not suppressed by guanine. Oasis 2 statistics for CLS assays are provided in Table S5.

We first considered the possibility that INAM was extending *S. cerevisiae* CLS by specifically perturbing guanine nucleotide pools through a mechanism different from MPA. As shown in Fig. 5D, MPA extended CLS at 1 µM, and even more effectively at 10 µM, consistent with its known extension of RLS (43) and *S. pombe* CLS (44). As predicted, CLS extension by MPA was fully reversed by supplementing with 0.1 mM guanine (Fig. 5D), which bypasses IMP dehydrogenase inhibition by conversion to GMP via the salvage enzyme hypoxanthine-guanine phosphoribosyltransferase (Hpt1; Fig. 5A). In contrast, CLS extension induced by 25 mM INAM was not reversed by guanine (Fig. 5E), implying that IMP dehydrogenase was not directly inhibited by INAM. Ura3 also cannot be a relevant INAM target because BY4741 and the YKO collection strains are all *ura3Δ0*.

### INAM perturbs nucleotide metabolism

The combination of INAM sensitivity for mutants defective in transcriptional elongation, de novo purine biosynthesis (*ADE* genes), or serine/threonine/glycine biosynthesis pathways, as well as synergistic growth inhibition with MPA suggested a broader effect on nucleotide metabolism. To test this idea, we utilized mass spectrometry to quantify a panel of nucleotides and precursors when BY4741 was grown in the presence of 25 mM INAM. Extracts were isolated from treated and untreated cells grown to log phase (6 hrs), late diauxic shift (24 hr), or stationary phase (96 hr). Several nucleosides and bases were significantly reduced by INAM in log phase cells (Fig. 6A). While INAM did not significantly affect nucleotide levels in log phase cells, there was still a trend toward reduction of NTPs and dNTPs that became more significant at the 24 hr and 96 hr timepoints (Fig. 6A and B). These changes were accompanied by reduced aspartate and glutamine, amino acids that contribute atoms to de novo biosynthesis of the purine and pyrimidine rings. UTP was the lone NTP/dNTP exception that was not significantly reduced by the 96 hr timepoint in INAM-treated cells (Fig. 6A). Moreover, uracil, uridine, UMP, and UDP were all strongly upregulated at 96 hr. These changes could be related to the *ura3Δ0* mutation in BY4741 that blocks de novo UMP synthesis, forcing the cells to salvage uracil from the media. We therefore tested whether supplementing extra uracil into the growth media would extend CLS and/or modify the lifespan extension induced by INAM. Adding uracil at 4x the normal concentration in SC media did significantly extend CLS but had little impact on lifespan extension induced by 25 mM INAM (Fig. 6C).

**Fig. 6.**
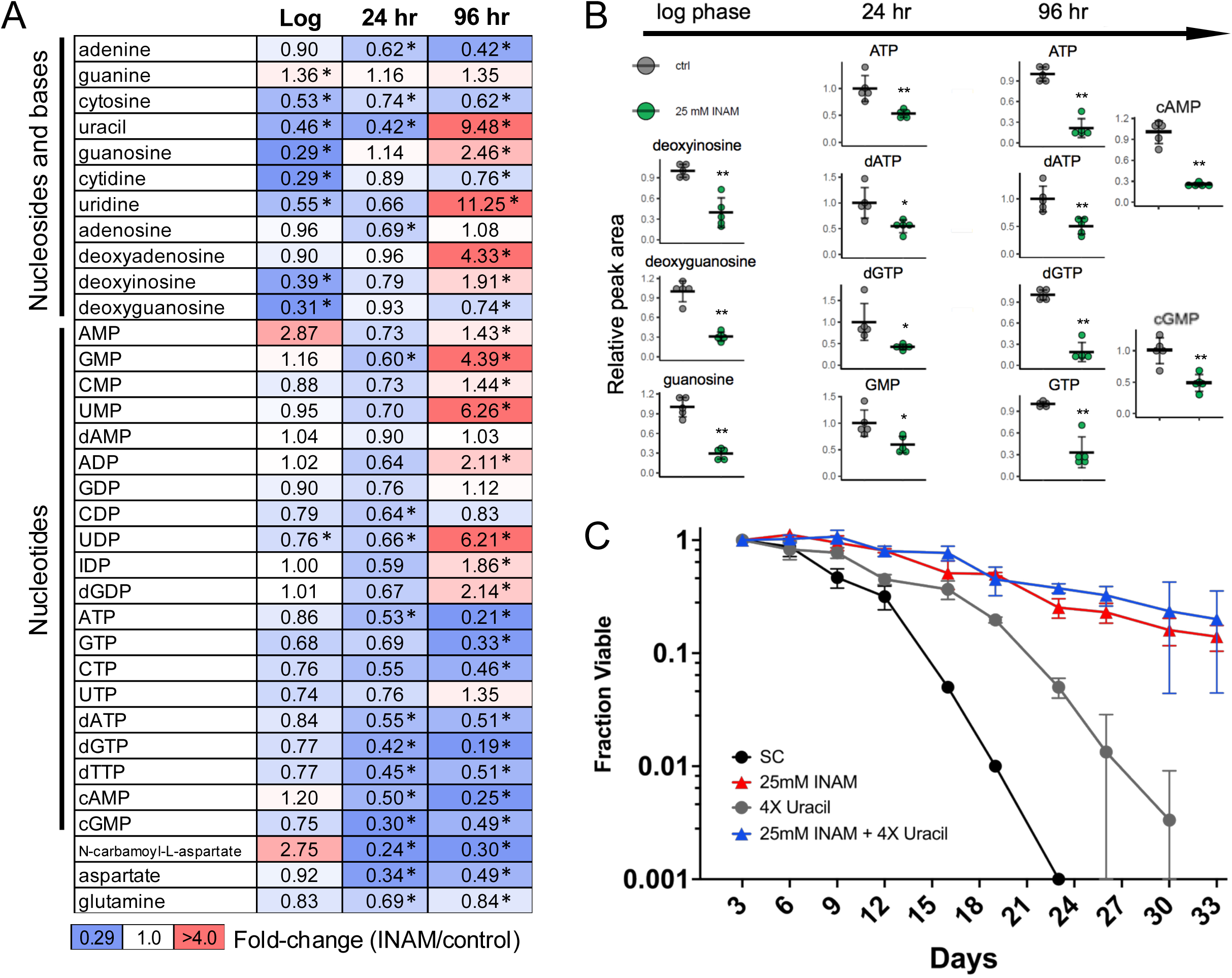
Alterations of intracellular nucleotide metabolite levels by 25 mM INAM. **A)** Summary of quantitative mass spectrometry analysis for a panel of purine and pyrimidine nucleotides, nucleosides, bases, and precursors. Block shading indicates fold-change relative to the control without INAM (red-up and blue-down). Asterisks indicate significant changes with p-values <0.05. **B)** Individual examples of significantly reduced purine nucleosides or nucleotides reduced by INAM at each time point. *p<0.05, **p<0.01. **C)** CLS assay showing partial lifespan extension when BY4741 (uracil auxotroph) is supplemented with 4x (0.8 mM) the normal concentration of uracil (0.2 mM) in SC medium. INAM (25 mM) fully extends CLS in combination with 4x uracil. Oasis 2 statistics for CLS assays are provided in Table S5.

Among the nucleotide-related metabolites significantly reduced by INAM during log phase were cytidine, guanosine, and uridine (Fig. 6A), nucleosides primarily produced by dephosphorylation of the ribonucleoside monophosphates, CMP, GMP, and UMP during nucleotide salvage (Fig. 7A). Sdt1 and Phm8 are paralogous 5’-NMP-specific nucleotidases (45, 46), and Isn1 is an IMP-specific 5’ nucleotidase (Fig. 7A). Additionally, the vacuolar alkaline phosphatase Pho8 dephosphorylates 3’-NMPs to generate cytidine, guanosine, and uridine during degradation of RNA by autophagy in response to nitrogen starvation (47). We hypothesized that the depletion of nucleosides caused by INAM was due to inhibition of nucleotidase and/or alkaline phosphatase activity. To test this, BY4741 cultures were grown into log phase, then INAM was added at concentrations of 25 or 100 mM for only 1 hr to track acute metabolite changes that were likely due to direct enzymatic inhibition. Cytidine and guanosine were again significantly reduced by INAM, and in a dose-dependent manner (Fig. 7B). While inosine was not included in these metabolite panels, its downstream base, hypoxanthine, was significantly reduced (Fig. 7B). Cytosine and uracil were similarly reduced by extended INAM exposure during log phase (Fig. 6A), suggesting downstream depletion of the bases, with the surprising exception of guanine.

**Fig. 7.**
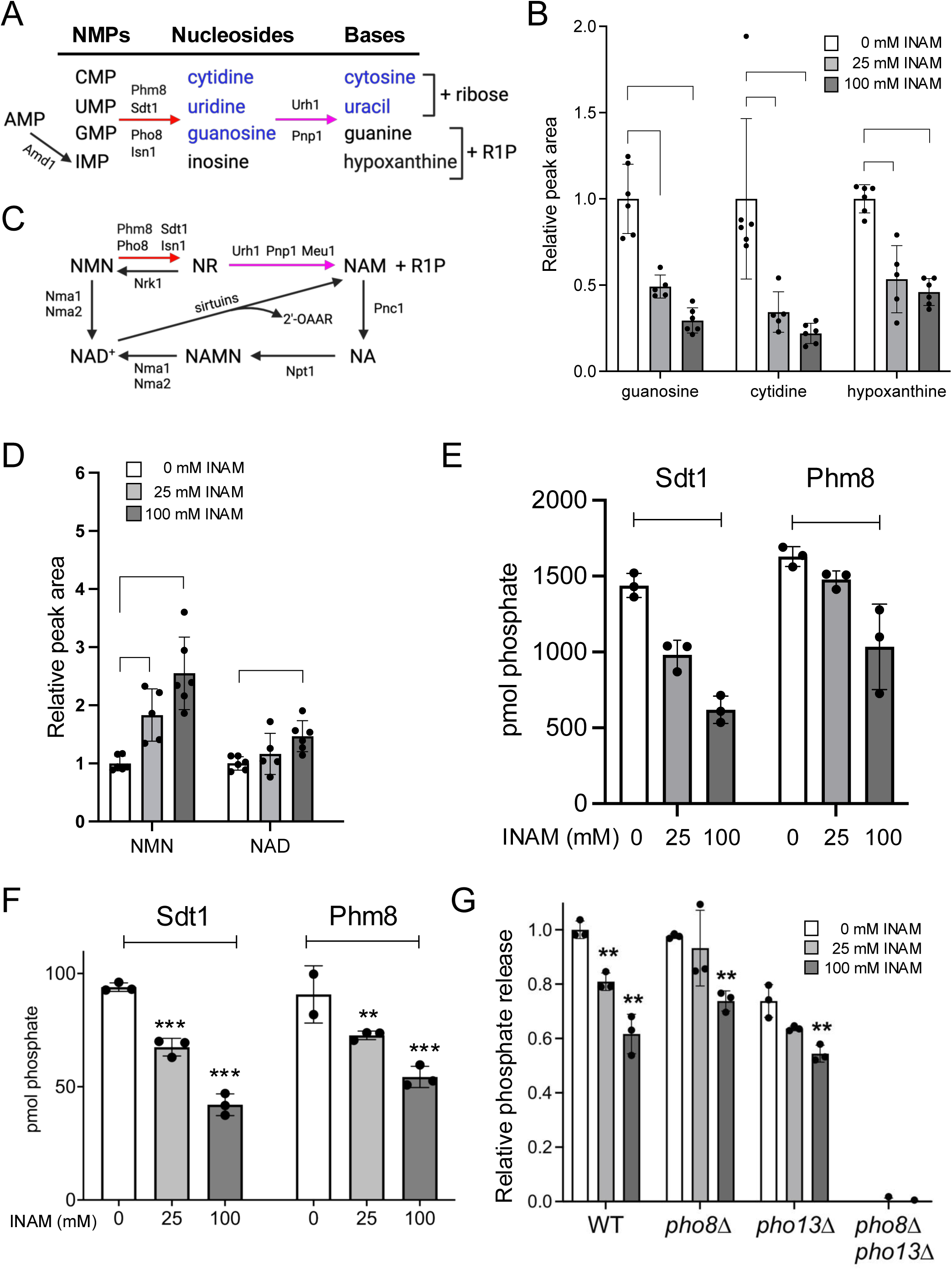
Inhibition of nucleotidase and alkaline phosphatase activity by INAM. **A)** Nucleoside monophosphate breakdown to bases by nucleotidases (Phm8, Sdt1, Isn1), alkaline phosphatases (Pho8) to produce nucleosides, followed by the action of purine nucleoside phosphorylase (Pnp1) and uridine nucleosidase (Urh1) to release the bases and either ribose or ribose-1-phosphate. **B)** Quantitation of guanosine, cytidine, and hypoxanthine levels following 1 hr treatment with 0, 25, or 100 mM INAM. The mean of the control samples without INAM was normalized to 1.0. **C)** Nicotinamide mononucleotide (NMN) is broken down to NAM by the same set of enzymes that degrade NMPs as part of the NAD salvage pathway. **D)** Quantitation of NMN and NAD^+^ levels by mass spectrometry following 1 hr treatment with 25 or 100 mM INAM. The control was normalized to 1.0. **E)** INAM inhibition of recombinant Sdt1 and Phm8 nucleotidase activity on CMP in vitro. **F)** INAM inhibition of recombinant Sdt1 nucleotidase activity on NMN in vitro. **G)** INAM inhibition of alkaline phosphatase activity in whole extracts isolated from WT, *pho8Δ*, *pho13Δ*, and *pho8Δ pho13Δ* strains using p-nitrophenyl phosphate (pNPP) as a substrate. In all graphs, error bars represent standard deviations. Significance p-values were *<0.05, **<0.005, ***<0.005, as calculated using 1-way Anova.

The same nucleotidases and phosphatases, Phm8, Sdt1, Isn1, and Pho8, have also been reported to convert NMN to NR (Fig. 7C; (48, 49)), which is intriguing given the beneficial effect of INAM on NAD^+^ homeostasis as cells enter the diauxic shift (14). NMN does not cross the yeast cell membrane, so it must first be dephosphorylated into NR and then imported by the NR transporter Nrt1 (50, 51). Preventing NMN dephosphorylation to NR by inhibition of Sdt1, Phm8, Isn1, and Pho8 could potentially ‘trap’ NMN in the cell and shift the flux toward NAD^+^ via nicotinamide adenylyltransferases Nma1/Nma1/Pof1 (Fig. 7C) (52, 53). We therefore measured changes in NMN and NAD^+^ when BY4741 was acutely treated with INAM for 1 hr. As shown in Fig. 7D, NMN accumulated in a dose-dependent manner that translated to elevated NAD^+^ at 100 mM INAM, also consistent with our hypothesis of nucleotidase/phosphatase inhibition.

### INAM inhibits nucleotidase and alkaline phosphatase activities

We next tested whether Sdt1 and Phm8 were direct targets of INAM *in vitro* by incubating the recombinant His-tagged enzymes with CMP or NMN substrates and quantifying the release of inorganic phosphate with a colorimetric malachite green phosphate detection assay (54). Sdt1 and Phm8 activity on CMP and NMN was significantly inhibited by INAM concentrations equivalent to those that impacted CLS and nucleoside levels in cultured cells (Figs. 7E-F). The effect of INAM on alkaline phosphatase activity was next tested by incubating p-nitrophenol phosphate (pNPP) with whole cell extracts from exponentially growing WT, *pho8Δ*, *pho13Δ*, and *pho8Δ pho13Δ* strains. Pho8 and cytosolic Pho13 are the only alkaline phosphatases in S. cerevisiae, and both utilize pNPP as a substrate in vitro (49, 55). Dephosphorylation activity on pNPP was eliminated in the *pho8Δ pho13Δ* double mutant extract (Fig. 7G), confirming specificity of the assay. INAM weakened alkaline phosphatase activity from the WT and single mutant extracts, suggesting that Pho8 and Pho13 were both weak inhibitory targets. Taken together, we conclude that INAM impairs a specific step of the nucleotide salvage pathway, conversion of NMPs to nucleosides. This likely accounts for the synthetic growth defects with transcriptional elongation mutants or pathways related to de novo nucleotide synthesis, including one-carbon metabolism inputs. These findings also provide a potential mechanism for the maintenance of NAD^+^ homeostasis by INAM without being directly metabolized.

## Discussion

### Isonicotinamide and NAD^+^ homeostasis

INAM originally came to our attention because it prevented feedback inhibition of Sir2 activity by NAM, a product of the lysine deacetylation reaction (13). Unexpectedly, it also promoted RLS in a Sir2-dependent manner by indirectly maintaining elevated NAD^+^ levels under conditions where NAD^+^ normally becomes depleted, such as media lacking nicotinic acid

(14). INAM also prevented NAD^+^ depletion as cells enter the diauxic shift (14), which led us to test for effects on CLS. INAM at 25 mM strongly extended CLS independently of Sir2 (15), and we now show that it also extends lifespan of a quintuple mutant strain lacking all sirtuins (Fig. 2A). While this result precludes Sir2 and other sirtuins mediating the beneficial effect of INAM on CLS, the improved NAD^+^ homeostasis could potentially be contributing to the CLS extension independent of sirtuins. Indeed, the NR salvage pathways (Nrk1 and Urh1/Pnp1/Meu1; Fig. 7C) were previously reported as critical for maintaining CLS (56). Our results suggest that INAM inhibition of the nucleotidases Sdt1 and Phm8, or alkaline phosphatases, shifts flux toward intracellular retention of NMN to promote NAD^+^ homeostasis, rather than the NMN being converted to NR.

### INAM effects compared to NAM

An earlier screen of the YKO collection for deletion mutants sensitive to nicotinamide (NAM) identified genes that function in sister chromatid cohesion and maintenance of genome stability (57), consistent with the increased mutation frequency caused by high NAM concentrations (Fig. 2C). From the supplemental tables of this paper (57) we were also able to extract 107 NAM-sensitive mutants with normalized colony growth ratios of <0.5 in both screen replicates (57), revealing significantly enriched GO terms of DNA repair (as expected), as well as de novo IMP biosynthesis (*ADE1*, *ADE2*, *ADE4*, *ADE6*, *ADE8*). Genome maintenance genes were not significantly enriched in our screen for INAM-sensitive mutants, consistent with the absence of any mutagenic effect. Interestingly, 21 of the 59 reported NAM-sensitive mutants did overlap with our list of INAM-sensitive mutants from Table S4, including *htz1Δ*. Notably, we identified several *ADE* gene mutants as INAM-sensitive, suggesting that NAM and INAM share some level of overlap in their metabolic effects on yeast cells, as might be expected for compounds with such similar chemical structures (Fig. 1A). While NAM at relatively low (1 or 5 mM) concentrations previously shortened CLS (23), most likely due to combined inhibition of Sir2 and Hst1, the extension induced by 25 mM appears to override any inhibitory effect on sirtuins, reminiscent of INAM extending CLS independently of the sirtuins.

### INAM effects on nucleotide metabolism

One of the key clues that INAM was potentially modifying nucleotide metabolism was identification of multiple INAM-sensitive elongation factor mutants that were also sensitive to MPA or 6-AU (39, 58). MPA and 6-AU reduce the guanine nucleotide pool by inhibiting IMPDH, and the combination of impaired elongation from the mutations and reduced nucleotide pools by these compounds caused growth defects in the previous studies. We therefore hypothesized that INAM was also inhibiting growth of isolated elongation factor mutants due to impairing nucleotide pools. Additional genetic support was derived from INAM-sensitive mutants in serine, threonine, aspartate, the transsulfuration pathway, and de novo purine biosynthesis (ADE) pathway, each of which can be linked to nucleotide synthesis via the 1C metabolism pathway. For these reasons we first directly measured nucleotide and precursor levels when cells were chronically grown in 25 mM INAM across a time course as they entered stationary phase. Nucleosides and bases in the test panel were significantly reduced in log phase cells (Fig. 6A), followed by significant reduction of dNTPs at 24 hr and then both NTPs and dNTPs at 96 hr. Acute INAM treatment of log phase cells for 1 hr confirmed the depletion of guanosine and cytidine, pointing to the nucleotidases Sdt1 and Phm8, or alkaline phosphate Pho8, as candidate targets for inhibition by INAM. We hypothesize that the INAM-induced limitation of several bases and nucleosides during log phase growth has a downstream effect of preventing efficient salvage synthesis of nucleotides and therefore accelerating the natural depletion of nucleotide pools as cells enter stationary phase (59). The lack of guanine depletion at any timepoint is surprising, yet consistent with guanine supplementation not influencing CLS in INAM treated cells. The amino acid precursors to de novo UMP synthesis, glutamine and aspartate, as well as the N-carbamoyl-L-aspartate intermediate were all depleted by INAM at 24 and 96 hrs, but since the BY4741 strain is deleted for *URA3* (OMP decarboxylase), it is likely that the elevated levels of uracil, uridine, UMP, and UDP at 96 hr are symptomatic of the blocked *de novo* pyrimidine synthesis. Lastly, given the structural similarity with NAM, we cannot rule out the possibility that INAM alters the activity of additional enzymatic steps of de novo or salvage pathways for nucleotide metabolism, or any enzyme that interacts with NAD(H) or NADP(H).

### Nucleotide pool effects on lifespan

In multicellular organisms, the rate of transcriptional elongation increases with age and genetic manipulations that slow elongation tend to extend lifespan (60). It remains unclear if alterations in elongation have any impact on yeast CLS. In our study, deletion of *SWR1* or *HTZ1* was not sufficient to extend CLS and did not block the CLS extension induced by 25 mM INAM, even though the *htz1Δ* mutant growth was slowed at this concentration. Therefore, any combined impacts on transcriptional elongation by these mutants and/or INAM were likely not driving the lifespan extension.

An alternative hypothesis is that the altered nucleotide pools post-log phase could be causing low level DNA replication stress, which was previously shown for guanine nucleotide depletion in T and B cells that causes S-phase arrest, replication stress, and apoptosis due to DNA replication fork stalling (61, 62). Hydroxyurea (HU) also reduces the dNTP pool by inhibiting ribonucleotide reductase to cause replication stress. Interestingly, HU extends CLS independent of activating various stress resistance pathways and instead enhances entry into quiescence (63). It was hypothesized that the replication stress would increase the proportion of cells transiently arresting the cell cycle during the growth phase, which would increase the likelihood of arresting cell growth and entering quiescence in subsequent cell cycles (63). Reducing the guanylic nucleotide pool with MPA was previously shown to cause a mother-daughter cell separation defect and emergence of bi-budded cells, which were observed in cells entering quiescence (64). Therefore, a common thread between HU and MPA is they appear to enhance the entry to quiescence when cultures approach stationary phase. Similarly, INAM induced DNA replication stress could be enhancing CLS through stimulating entry into quiescence.

### cAMP and chronological aging

Additional evidence for improved quiescence entry by INAM supplementation is derived from the significant reduction of cAMP and cGMP at the 24 and 96 hr time points. cAMP levels are typically high in proliferating yeast cells and then reduced as cells enter stationary phase, controlled by the balance between the activities of adenylyl cyclase and the cAMP phosphodiesterases, Pde1 and Pde2 (65). The reduction in cAMP/PKA signalling allows Rim15 to enter the nucleus and activate genes with Post-Diauxic Shift (PDS) or STress Response Element (STRE) motifs. This cascade is key to the mechanism of glucose restriction mediated RLS extension (66). Consistent with this idea, forcing reduction in cAMP by inhibiting adenylate cyclase (Cyr1) with the drug triclabendazole has been shown to extend yeast CLS (67). The observed cAMP reduction in INAM-treated cells at 24 and 96 hr is consistent with lifespan extension and suggests possible stimulation of Pde1/Pde2 activity and/or reduced Cyr1 activity. Alternatively, and more likely, is that the reduced cAMP levels are linked to reduced ATP, the substrate for Cyr1. Either way, the reduction in cAMP caused by INAM could be enhancing stationary phase entry and quiescence maintenance via Rim15 activation, thus contributing to CLS extension. Little is known about the role of cGMP in yeast cells, but it has been reported to activate PKA (68), and proposed to help fine tune cAMP levels (69), so it could also potentially play a role in INAM-mediated CLS extension.

### Isonicotinamide implications for human health

Could INAM or its derivatives be used as an aging intervention in humans? INAM is commonly used as a backbone for the synthesis of various drugs, as well as cocrystallization with existing compounds to make them more bioavailable (70, 71). Examples include the MEK1/2 inhibitor pimasertib (72), xanthine oxidase inhibitors where the INAM moiety is the key functional region (73), and new GSK-3 inhibitors in development as Alzheimer’s treatments (74). INAM itself is also known to inhibit the activity of PARP-1 in mammalian cells, similar to the effect of NAM (75). Indeed, the strong elevation of NAD^+^ observed in lung cancer cells upon INAM treatment is likely caused by PARP-1 inhibition, though nucleotidases/phosphatase inhibition could also be involved. Given its structural similarity to NAM, INAM also has the potential to target other enzymes that interact with the NAD^+^ family of metabolites. As INAM-derived therapeutics are used more in clinical trials, it will therefore be of interest to track the impacts on NAD^+^ levels and nucleotide pools, as well as healthspan parameters.

## MATERIALS and METHODS

### Yeast strains and media

The primary ‘wild-type’ laboratory strain in this study was BY4741 (*MAT*a *his3Δ1 leu2Δ0 met15Δ0 ura3Δ0*) (76), and most gene deletion mutants were obtained from the isogenic yeast knockout (YKO) collection (77). Detailed strain information is provided in Supplemental Table S4. The *ade2-101* mutation in YPH499 and YCB498 was repaired by transformation with an *ADE2* PCR product spanning the ochre mutation and selecting for Ade^+^ colonies to yield SY1043 and SY1044, respectively. The reversion was confirmed by Sanger DNA sequencing. Synthetic complete (SC) media with 2% glucose as a carbon source was used for most assays, with YPD used for cell propagation and selection prior to the experiments. All liquid cultures and agar plates were grown at 30°C.

### CLS assays

10 ml of SC media with 2% glucose and relevant supplements was inoculated with 100 µl of overnight culture. Liquid cultures were incubated on a rotating drum (TC-7, New Brunswick Scientific) in glass tubes with metal caps allowing for gas exchange. After 72 hours, the first measurement of colony forming units on YPD agar plates was made and this was treated as day 3 for the experiment (100% starting viability), to which all the other data was normalized. Measurements were taken every 3-4 days as previously described (29, 78). At each time point, 20 µl of the cell suspension was removed from each tube and 10-fold serially diluted three times with sterile water. Next, 2.5 µl of each dilution (1:10, 1:100, 1:1000) was spotted onto a YPD plate. After 16 hours, images of the spots were taken under a Nikon Eclipse E400 brightfield microscope at 30x magnification. Microcolonies within the spots were automatically counted from the digital images using OrganoSeg (79), with the parameters adjusted for yeast colony counting (29). After accounting for the dilution factor, colony numbers from each day were divided by the number of colonies from the first time point (day 3) to give the viability metrics.

### SGA screen

The *MAT*a haploid YKO collection, consisting of 4648 strains organized into 18 plates in 384-well format, was spotted onto Nunc Omnitray single-well plates containing YPD-agar with 200 µg/ml G418 using a floating pin manual replicator (VP 384FS, V&P Scientific). After 3 days growth, the replicator was used to transfer cells from colonies on the YPD plates to SC-agar Omnitrays supplemented with 0, 25, 50, 75, or 125 mM INAM. Colony growth for the 0, 25, 50, and 75 mM INAM plates was imaged with a Fluor Chem Q gel documentation system (ProteinSimple) after 2 days, while the 125 mM plates were imaged after 3 days incubation. The screen was performed in duplicate. SGAtools (http://sgatools.ccbr.utoronto.ca/) was then used to quantify colony growth from jpeg images, comparing the INAM-containing plates to control plates without INAM (27). The score value of -0.3 was used as a cutoff for INAM sensitivity, and to limit false positives we required that both replicates reach this threshold for at least the 75 and 125 mM conditions. Most mutants with deletions of dubious open reading frames and genes without a confirmed function were omitted in further analysis, as their sensitivity would not provide any valuable information about the mechanism of action of INAM.

### Growth assays

For qualitative spots test growth assays, strains were patched onto YPD agar plates and grown overnight. The cell patches were resuspended in water at an OD_600_ of 1.0, then 10-fold serially diluted. Next, 2.5µl of each dilution was spotted onto SC agar plates containing 0, 25, 50, 75, or 125 mM INAM, and photos taken after 2 days of growth.

For IC_50_ assays, 10 ml SC cultures were incubated overnight in the roller drum, then added to wells of a 96-well plate at a starting OD_600_ of 0.1 in SC media containing a range of INAM concentrations (0-240 mM; 2-fold dilutions) and other supplements as indicated. The initial OD_600_ was measured with a Spectra Max M2 plate reader (Molecular Devices). Plates were then sealed with a sterile, gas-permeable membrane and incubated in a heated microplate shaker (Southwest Science, SBT1500-H) at 30°C for 8 hr at 900 RPM. Final OD_600_ was measured and the initial OD_600_ subtracted. Values were then normalized to fit the range 0-100%, where the control condition (no INAM) was treated as 100% and the highest INAM concentration as 0. These normalized growth values were then plotted on a graph using Origin 2018 (OriginLab, Northampton, MA, USA) and the sigmoidal fit was used to model a dose-response curve fitting the obtained data. IC_50_ was calculated as the concentration at which there was 50% viability. The 95% confidence intervals were also calculated by the Origin software from the 95% confidence bands. An additional benefit of this IC_50_ approach is precisely determining if mutant strains with intrinsically slow growth are sensitive to INAM, due to the assay being internally normalized to growth of a given strain in the control condition (no INAM), thus allowing direct comparison of strains with different growth rates.

Synergy assessment was performed similarly to the IC_50_ assays, with a combination of increasing MPA and INAM concentrations added in a matrix to a 96-well plate. Methanol vehicle concentration was adjusted to be the same at all MPA concentrations used. SynergyFinder (80) was used to analyze the synergy between compounds using baseline correction and a zero interaction potency (ZIP) model of synergy determination (81).

### Measurement of metabolite abundance by mass spectrometry

Cells were grown in 10 ml SC 2% glucose with or without 25 mM INAM on a roller drum. Cultures were started from overnight liquid cultures at OD_600_ adjusted to 0.05. Cell densities of cultures were measured before collection for normalization of metabolites. Cells were collected at log phase (∼5 hr), 24 hr, and 96 hr. Alternatively, log phase cultures were supplemented with 0, 25, or 100 mM INAM for 1 hr, followed by immediate cell harvest. Intracellular metabolites were isolated as previously described (82), with the extraction protocol modified for yeast. From these cultures, 9 ml was added to 20 ml of 100% cold (-80°C) LC/MS-grade methanol kept on dry ice (final methanol concentration: ∼70% v/v) (Canelas *et al*, 2008). Samples were then kept on dry ice unless indicated otherwise. The quenched cell suspensions were spun down at 2000 × g in an Eppendorf 5810R centrifuge cooled to -9°C for 5 min. The cell pellets were resuspended in 1 ml 80% methanol (LC/MS grade) and transferred to screw-top microfuge tubes containing 0.5 mm acid-washed glass beads. A Mini-Beadbeater (Biospec products) was used to vigorously shake the microfuge tubes at 4°C for 3 cycles of 45 sec with 30-second pauses. Each tube was then punctured at the bottom with a needle, placed into a glass tube and spun down at -9°C for 5 minutes at 2000 RPM in the Eppendorf 5810R centrifuge to collect cell extracts. Beads were washed twice with 1.5 ml 80% methanol and spun down the same way (4 ml total pooled volume). The pellet was resuspended by vortexing and kept on ice for 15 minutes, followed by centrifugation at 3100 × g. Supernatants were transferred to two 2-ml microfuge tubes, and evaporated in a Vacufuge™ Concentrator 5301 (Eppendorf, Hamburg, Germany) at 30°C until 20% volume remained. Remaining water was then removed by lyophilization.

Metabolites were analyzed as described previously (83, 84), with a selected reaction monitoring (SRM) LC-MS/MS method with positive/negative ion polarity switching using a Xevo TQ-S mass spectrometer. MassLynx 4.1 software was used to calculate the peak areas for each metabolite. Peak areas were normalized to respective OD_600_ of each sample. These normalized peaks were subsequently scaled relative to the mean peak area in the control condition at each time point.

### Enzymatic assays

Alkaline phosphatase activity was measured from whole cell extracts as previously described (49). WT (BY4741), *pho8Δ* (SY1124), *pho13Δ* (SY1136) and *pho8Δ pho13Δ* (SY1137) strains were grown overnight in 10 ml YPD to an A_600_ of 2 to 3. Cell pellets were washed (0.85% NaCl, 1 mM PMSF) and then disrupted in 1.8 ml lysis buffer (20mM PIPES, 0.5% Triton X-100, 50mM KCl, 100mM potassium acetate, 10mM MgSO_4_, and 10µM ZnSO_4_ with 1mM PMSF added just before use) using a Biospec Mini-Beadbeater. Following centrifugation, 200µl of supernatant was added to 300µl of reaction buffer (333mM Tris-HCl, pH 8.8, 133mM MgSO_4_, 13.3µM ZnSO_4_, 0.53% Triton X-100, with or without 1.66mM p-Nitrophenyl Phosphate) in a microfuge tube and incubated for 15 min at 37°C. Next, 500µl of stop buffer (1M Glycine, 1M KOH, pH 11) was added to the reaction tubes, which were centrifuged again for 5 min at 4°C. 200µl of each reaction was then transferred into a 96-well plate and read at A_400_ on a Molecular Devices Spectramax M5 plate reader. Activity was normalized to the A_600_ of each culture.

Nucleotidase activity of recombinant Sdt1 and Phm8 was measured by quantifying the release of phosphate from in vitro reactions. C-terminally His-tagged Sdt1 (E-20SDT1) was obtained from Echelon Biosciences Inc. (Salt Lake City, Utah). Phm8 was cloned into the *Nde*I site of pET-16b plasmid and expressed in BL21(DE3) *E. coli* as a C-terminal 10xHis-tagged protein. The recombinant protein was purified using Ni-NTA agarose beads and the pooled fractions were dialyzed with 50 mM Tris-HCl, pH 7.5, 50 mM NaCl, 2 mM β-mercaptoethanol, 50% glycerol in 30 kDa cutoff dialysis tubing. The Sigma-Aldrich Phosphate Assay Kit (MAK308-1KT) was used to determine nucleotidase activity. 0.5 μg Sdt1 or 0.5 μg Phm8 was incubated with 100 µM CMP (Tokyo Chemical Industry, YK6JM-MI) or NMN (Cayman, 16411) for 30 min at 37°C in 50 µl of reaction buffer (50 mM Tris-HCl, pH 7.5, 5 mM MgCl_2_) in a 96-well plate. CMP and NMN were left out as negative controls. Reactions were then incubated with 100 µl of malachite green solution (Sigma, MAK308A-KC) for minutes at room temperature (54). Absorbance was then measured at 620 nm using the Spectramax M5 plate reader. Dilutions of a 1 mM phosphate standard were used as a standard curve for quantitation of pmol phosphate released.

## Supporting information

This article contains supporting information

## Acknowledgments

We thank members of the Smith lab, David Auble, Dan Burke, and Marty Mayo for helpful discussions, as well as Dan Burke, Jef Boeke, Phil Hieter, Mitch Smith, and Fred Winston for kindly providing yeast strains. E.E.H was supported by the Medical Scientist Training Program (MSTP) NIH training grant T32GM007267 and an individual National Research Service Award (NRSA) F30AG067760. L.P.N. was supported by NRSA F31AG081044. Cell and Molecular Biology (CMB) NIH training grant T32GM008136 partially supported E.E.H. and C.T.L. The study was also supported by NIH grants R01GM075240 and R01GM127394 to J.S.S., as well as R01CA210439 and R01CA163649 to P.K.S.

## References

1. Bitto, A., Wang, A. M., Bennett, C. F., and Kaeberlein, M. (2015) Biochemical Genetic Pathways that Modulate Aging in Multiple Species. Cold Spring Harb Perspect Med 5, a025114

2. Smith, E. D., Tsuchiya, M., Fox, L. A., Dang, N., Hu, D., Kerr, E. O. et al. (2008) Quantitative evidence for conserved longevity pathways between divergent eukaryotic species. Genome Res 18, 564–570

3. Gartenberg, M. R., and Smith, J. S. (2016) The nuts and bolts of transcriptionally silent chromatin in *Saccharomyces cerevisiae*. Genetics 203, 1563–1599

4. Brachmann, C. B., Sherman, J. M., Devine, S. E., Cameron, E. E., Pillus, L., and Boeke, J. D. (1995) The *SIR2* gene family, conserved from bacteria to humans, functions in silencing, cell cycle progression, and chromosome stability. Genes Dev 9, 2888–2902

5. Kaeberlein, M., McVey, M., and Guarente, L. (1999) The SIR2/3/4 complex and SIR2 alone promote longevity in *Saccharomyces cerevisiae* by two different mechanisms. Genes Dev 13, 2570–2580

6. Lin, Y. Y., Lu, J. Y., Zhang, J., Walter, W., Dang, W., Wan, J. et al. (2009) Protein acetylation microarray reveals that NuA4 controls key metabolic target regulating gluconeogenesis. Cell 136, 1073–1084

7. Casatta, N., Porro, A., Orlandi, I., Brambilla, L., and Vai, M. (2013) Lack of Sir2 increases acetate consumption and decreases extracellular pro-aging factors. Biochim Biophys Acta 1833, 593–601

8. Anderson, R. M., Bitterman, K. J., Wood, J. G., Medvedik, O., and Sinclair, D. A. (2003) Nicotinamide and PNC1 govern lifespan extension by calorie restriction in *Saccharomyces cerevisiae*. Nature 423, 181–185

9. Bitterman, K. J., Anderson, R. M., Cohen, H. Y., Latorre-Esteves, M., and Sinclair, D. A. (2002) Inhibition of silencing and accelerated aging by nicotinamide, a putative negative regulator of yeast Sir2 and human SIRT1. J Biol Chem 277, 45099–45107

10. Gallo, C. M., Smith, D. L., Jr., and Smith, J. S. (2004) Nicotinamide clearance by Pnc1 directly regulates Sir2-mediated silencing and longevity. Mol Cell Biol 24, 1301–1312

11. Orlandi, I., Pellegrino Coppola, D., Strippoli, M., Ronzulli, R., and Vai, M. (2017) Nicotinamide supplementation phenocopies SIR2 inactivation by modulating carbon metabolism and respiration during yeast chronological aging. Mech Ageing Dev 161, 277–287

12. Sauve, A. A., and Schramm, V. L. (2003) Sir2 regulation by nicotinamide results from switching between base exchange and deacetylation chemistry. Biochemistry 42, 9249–9256

13. Sauve, A. A., Moir, R. D., Schramm, V. L., and Willis, I. M. (2005) Chemical activation of Sir2-dependent silencing by relief of nicotinamide inhibition. Mol Cell 17, 595–601

14. McClure, J. M., Wierman, M. B., Maqani, N., and Smith, J. S. (2012) Isonicotinamide enhances Sir2 protein-mediated silencing and longevity in yeast by raising intracellular NAD^+^ concentration. J Biol Chem 287, 20957–20966

15. Wierman, M. B., Matecic, M., Valsakumar, V., Li, M., Smith, D. L., Jr., Bekiranov, S. et al. (2015) Functional genomic analysis reveals overlapping and distinct features of chronologically long-lived yeast populations. Aging 7, 177–194

16. Fleming, M. A., Chambers, S. P., Connelly, P. R., Nimmesgern, E., Fox, T., Bruzzese, F. J. et al. (1996) Inhibition of IMPDH by mycophenolic acid: dissection of forward and reverse pathways using capillary electrophoresis. Biochemistry 35, 6990–6997

17. Mitsui, A., and Suzuki, S. (1969) Immunosuppressive effect of mycophenolic acid. J Antibiot 22, 358–363

18. Winston, F., Dollard, C., and Ricupero-Hovasse, S. L. (1995) Construction of a set of convenient *Saccharomyces cerevisiae* strains that are isogenic to S288C. Yeast 11, 53–55

19. Belenky, P., Racette, F. G., Bogan, K. L., McClure, J. M., Smith, J. S., and Brenner, C. (2007) Nicotinamide riboside promotes Sir2 silencing and extends lifespan via Nrk and Urh1/Pnp1/Meu1 pathways to NAD^+^. Cell 129, 473–484

20. Fine, R. D., Maqani, N., Li, M., Franck, E., and Smith, J. S. (2019) Depletion of limiting rDNA structural complexes triggers chromosomal instability and replicative aging of *Saccharomyces cerevisiae*. Genetics 212, 75–91

21. Hickman, M. A., and Rusche, L. N. (2007) Substitution as a mechanism for genetic robustness: the duplicated deacetylases Hst1p and Sir2p in S*accharomyces cerevisiae*. PLoS Genet 3, e126

22. Li, M., Valsakumar, V., Poorey, K., Bekiranov, S., and Smith, J. S. (2013) Genome-wide analysis of functional sirtuin chromatin targets in yeast. Genome Biol 14, R48

23. Smith, D. L., Jr., McClure, J. M., Matecic, M., and Smith, J. S. (2007) Calorie restriction extends the chronological lifespan of *Saccharomyces cerevisiae* independently of the Sirtuins. Aging Cell 6, 649–662

24. Tsuchiya, M., Dang, N., Kerr, E. O., Hu, D., Steffen, K. K., Oakes, J. A. et al. (2006) Sirtuin-independent effects of nicotinamide on lifespan extension from calorie restriction in yeast. Aging Cell 5, 505–514

25. Celic, I., Masumoto, H., Griffith, W. P., Meluh, P., Cotter, R. J., Boeke, J. D. et al. (2006) The sirtuins Hst3 and Hst4p preserve genome integrity by controlling histone H3 lysine 56 deacetylation. Curr Biol 16, 1280–1289

26. Kadyrova, L. Y., Mertz, T. M., Zhang, Y., Northam, M. R., Sheng, Z., Lobachev, K. S. et al. (2013) A reversible histone H3 acetylation cooperates with mismatch repair and replicative polymerases in maintaining genome stability. PLoS Genet 9, e1003899

27. Wagih, O., Usaj, M., Baryshnikova, A., VanderSluis, B., Kuzmin, E., Costanzo, M. et al. (2013) SGAtools: one-stop analysis and visualization of array-based genetic interaction screens. Nucleic Acids Res 41, W591–596

28. Ducker, G. S., and Rabinowitz, J. D. (2017) One-carbon metabolism in health and disease. Cell Metab 25, 27–42

29. Enriquez-Hesles, E., Smith, D. L., Jr., Maqani, N., Wierman, M. B., Sutcliffe, M. D., Fine, R. D. et al. (2021) A cell-nonautonomous mechanism of yeast chronological aging regulated by caloric restriction and one-carbon metabolism. J Biol Chem 296, 100125

30. Maruyama, Y., Ito, T., Kodama, H., and Matsuura, A. (2016) Availability of amino acids extends chronological lifespan by suppressing hyper-acidification of the environment in *Saccharomyces cerevisiae*. PLoS One 11, e0151894

31. Jung, P. P., Zhang, Z., Paczia, N., Jaeger, C., Ignac, T., May, P., et al. (2018) Natural variation of chronological aging in the *Saccharomyces cerevisiae* species reveals diet-dependent mechanisms of life span control. NPJ Aging Mech Dis 4, 3

32. Aon, M. A., Bernier, M., Mitchell, S. J., Di Germanio, C., Mattison, J. A., Ehrlich, M. R., et al. (2020) Untangling determinants of enhanced health and lifespan through a multi-omics approach in mice. Cell Metab 32, 100–116.e104

33. Pan, C., Li, B., and Simon, M. C. (2021) Moonlighting functions of metabolic enzymes and metabolites in cancer. Mol Cell 81, 3760–3774

34. Mizuguchi, G., Shen, X., Landry, J., Wu, W. H., Sen, S., and Wu, C. (2004) ATP-driven exchange of histone H2AZ variant catalyzed by SWR1 chromatin remodeling complex. Science 303, 343–348

35. Santisteban, M. S., Kalashnikova, T., and Smith, M. M. (2000) Histone H2A.Z regulates transcription and is partially redundant with nucleosome remodeling complexes. Cell 103, 411–422

36. Santisteban, M. S., Hang, M., and Smith, M. M. (2011) Histone variant H2A.Z and RNA polymerase II transcription elongation. Mol Cell Biol 31, 1848–1860

37. Wu, W. H., Wu, C. H., Ladurner, A., Mizuguchi, G., Wei, D., Xiao, H. et al. (2009) N terminus of Swr1 binds to histone H2AZ and provides a platform for subunit assembly in the chromatin remodeling complex. J Biol Chem 284, 6200–6207

38. Garay, E., Campos, S. E., González de la Cruz, J., Gaspar, A. P., Jinich, A., and Deluna, A. (2014) High-resolution profiling of stationary-phase survival reveals yeast longevity factors and their genetic interactions. PLoS Genet 10, e1004168

39. Desmoucelles, C., Pinson, B., Saint-Marc, C., and Daignan-Fornier, B. (2002) Screening the yeast “disruptome” for mutants affecting resistance to the immunosuppressive drug, mycophenolic acid. J Biol Chem 277, 27036–27044

40. Exinger, F., and Lacroute, F. (1992) 6-Azauracil inhibition of GTP biosynthesis in *Saccharomyces cerevisiae*. Curr Genet 22, 9–11

41. Escobar-Henriques, M., Balguerie, A., Monribot, C., Boucherie, H., and Daignan-Fornier, B. (2001) Proteome analysis and morphological studies reveal multiple effects of the immunosuppressive drug mycophenolic acid specifically resulting from guanylic nucleotide depletion. J Biol Chem 276, 46237–46242

42. Qiu, Y., Fairbanks, L. D., Rückermann, K., Hawrlowicz, C. M., Richards, D. F., Kirschbaum, B. et al. (2000) Mycophenolic acid-induced GTP depletion also affects ATP and pyrimidine synthesis in mitogen-stimulated primary human T-lymphocytes. Transplantation 69, 890–897

43. Liu, P., Sarnoski, E. A., Olmez, T. T., Young, T. Z., and Acar, M. (2020) Characterization of the impact of GMP/GDP synthesis inhibition on replicative lifespan extension in yeast. Curr Genet 66, 813–822

44. Stephan, J., Franke, J., and Ehrenhofer-Murray, A. E. (2013) Chemical genetic screen in fission yeast reveals roles for vacuolar acidification, mitochondrial fission, and cellular GMP levels in lifespan extension. Aging Cell 12, 574–583

45. Nakanishi, T., and Sekimizu, K. (2002) SDT1/SSM1, a multicopy suppressor of S-II null mutant, encodes a novel pyrimidine 5’-nucleotidase. J Biol Chem 277, 22103–22106

46. Xu, Y. F., Létisse, F., Absalan, F., Lu, W., Kuznetsova, E., Brown, G. et al. (2013) Nucleotide degradation and ribose salvage in yeast. Mol Syst Biol 9, 665

47. Huang, H., Kawamata, T., Horie, T., Tsugawa, H., Nakayama, Y., Ohsumi, Y. et al. (2015) Bulk RNA degradation by nitrogen starvation-induced autophagy in yeast. EMBO J 34, 154–168

48. Bogan, K. L., Evans, C., Belenky, P., Song, P., Burant, C. F., Kennedy, R. et al. (2009) Identification of Isn1 and Sdt1 as glucose-and vitamin-regulated nicotinamide mononucleotide and nicotinic acid mononucleotide 5’-nucleotidases responsible for production of nicotinamide riboside and nicotinic acid riboside. J Biol Chem 284, 34861–34869

49. Lu, S. P., and Lin, S. J. (2011) Phosphate-responsive signaling pathway is a novel component of NAD^+^ metabolism in *Saccharomyces cerevisiae*. J Biol Chem 286, 14271–14281

50. Belenky, P. A., Moga, T. G., and Brenner, C. (2008) *Saccharomyces cerevisiae YOR071C* encodes the high affinity nicotinamide riboside transporter Nrt1. J Biol Chem 283, 8075–8079

51. Lu, J. Y., Lin, Y. Y., Sheu, J. C., Wu, J. T., Lee, F. J., Chen, Y. et al. (2011) Acetylation of yeast AMPK controls intrinsic aging independently of caloric restriction. Cell 146, 969–979

52. Anderson, R. M., Bitterman, K. J., Wood, J. G., Medvedik, O., Cohen, H., Lin, S. S. et al. (2002) Manipulation of a nuclear NAD^+^ salvage pathway delays aging without altering steady-state NAD^+^ levels. J Biol Chem 277, 18881–18890

53. Kato, M., and Lin, S. J. (2014) YCL047C/POF1 is a novel nicotinamide mononucleotide adenylyltransferase (NMNAT) in *Saccharomyces cerevisiae*. J Biol Chem 289, 15577–15587

54. Lanzetta, P. A., Alvarez, L. J., Reinach, P. S., and Candia, O. A. (1979) An improved assay for nanomole amounts of inorganic phosphate. Anal Biochem 100, 95–97

55. Kaneko, Y., Toh-e, A., Banno, I., and Oshima, Y. (1989) Molecular characterization of a specific p-nitrophenylphosphatase gene, PHO13, and its mapping by chromosome fragmentation in *Saccharomyces cerevisiae*. *Mol Gen Genet* **220**, 133-139

56. Lu, S. P., Kato, M., and Lin, S. J. (2009) Assimilation of endogenous nicotinamide riboside is essential for calorie restriction-mediated life span extension in *Saccharomyces cerevisiae*. J Biol Chem 284, 17110–17119

57. Choy, J. S., Qadri, B., Henry, L., Shroff, K., Bifarin, O., and Basrai, M. A. (2015) A genome-wide screen with nicotinamide to identify sirtuin-dependent pathways in *Saccharomyces cerevisiae*. G3 **6**, 485–494

58. Lennon, J. C., 3rd, Wind, M., Saunders, L., Hock, M. B., and Reines, D. (1998) Mutations in RNA polymerase II and elongation factor SII severely reduce mRNA levels in *Saccharomyces cerevisiae*. *Mol Cell Biol* **18**, 5771-5779

59. Osório, H., Silles, E., Maia, R., Peleteiro, B., Moradas-Ferreira, P., Günther Sillero, M. A. et al. (2005) Influence of chronological aging on the survival and nucleotide content of *Saccharomyces cerevisiae* cells grown in different conditions: occurrence of a high concentration of UDP-N-acetylglucosamine in stationary cells grown in 2% glucose. FEMS Yeast Res 5, 387–398

60. Debès, C., Papadakis, A., Grönke, S., Karalay, Ö., Tain, L. S., Mizi, A. et al. (2023) Ageing-associated changes in transcriptional elongation influence longevity. Nature 616, 814–821

61. Laliberté, J., Yee, A., Xiong, Y., and Mitchell, B. S. (1998) Effects of guanine nucleotide depletion on cell cycle progression in human T lymphocytes. Blood 91, 2896–2904

62. Saxena, S., and Zou, L. (2022) Hallmarks of DNA replication stress. Mol Cell 82, 2298–2314

63. Ross, E. M., and Maxwell, P. H. (2018) Low doses of DNA damaging agents extend *Saccharomyces cerevisiae* chronological lifespan by promoting entry into quiescence. Exp Gerontol 108, 189–200

64. Sagot, I., Schaeffer, J., and Daignan-Fornier, B. (2005) Guanylic nucleotide starvation affects *Saccharomyces cerevisiae* mother-daughter separation and may be a signal for entry into quiescence. BMC Cell Biol 6, 24

65. Park, J. I., Grant, C. M., and Dawes, I. W. (2005) The high-affinity cAMP phosphodiesterase of Saccharomyces cerevisiae is the major determinant of cAMP levels in stationary phase: involvement of different branches of the Ras-cyclic AMP pathway in stress responses. Biochem Biophys Res Commun 327, 311–319

66. Wei, M., Fabrizio, P., Hu, J., Ge, H., Cheng, C., Li, L. et al. (2008) Life span extension by calorie restriction depends on Rim15 and transcription factors downstream of Ras/PKA, Tor, and Sch9. *PLoS Genet* **4**, e13

67. Lee, Y. J., Shi, R., and Witt, S. N. (2013) The small molecule triclabendazole decreases the intracellular level of cyclic AMP and increases resistance to stress in *Saccharomyces cerevisiae*. PLoS One 8, e64337

68. Cytryńska, M., Wojda, I., Frajnt, M., and Jakubowicz, T. (1999) PKA from *Saccharomyces cerevisiae* can be activated by cyclic AMP and cyclic GMP. Can J Microbiol 45, 31–37

69. Cardarelli, S., Giorgi, M., Poiana, G., Biagioni, S., and Saliola, M. (2019) Metabolic role of cGMP in *S. cerevisiae*: the murine phosphodiesterase-5 activity affects yeast cell proliferation by altering the cAMP/cGMP equilibrium. FEMS Yeast Res 19,

70. Báthori, N. B., Lemmerer, A., Venter, G. A., Bourne, S. A., and Caira, M. R. (2011) Pharmaceutical co-crystals with isonicotinamide—vitamin B3, clofibric acid, and diclofenac— and two Isonicotinamide hydrates. Crystal Growth & Design 11, 75–87

71. Sánchez-Férez, F., Ejarque, D., Calvet, T., Font-Bardia, M., and Pons, J. (2019) Isonicotinamide-based compounds: from cocrystal to polymer. Molecules 24, 4169

72. Akinleye, A., Furqan, M., Mukhi, N., Ravella, P., and Liu, D. (2013) MEK and the inhibitors: from bench to bedside. J Hematol Oncol 6, 27

73. Zhang, T. J., Li, S. Y., Wang, L., Sun, Q., Wu, Q. X., Zhang, Y. et al. (2017) Design, synthesis and biological evaluation of N-(4-alkoxy-3-cyanophenyl)isonicotinamide/nicotinamide derivatives as novel xanthine oxidase inhibitors Eur J Med Chem 141, 362–372

74. Luo, G., Chen, L., Burton, C. R., Xiao, H., Sivaprakasam, P., Krause, C. M. et al. (2016) Discovery of isonicotinamides as highly selective, brain penetrable, and orally active glycogen synthase kinase-3 Inhibitors. J Med Chem 59, 1041–1051

75. Ueda, K., Banasik, M., Nakajima, S., Yook, H. Y., and Kido, T. (1995) Cell differentiation induced by poly(ADP-ribose) synthetase inhibitors Biochimie 77, 368–373

76. Brachmann, C. B., Davies, A., Cost, G. J., Caputo, E., Li, J., Hieter, P. et al. (1998) Designer deletion strains derived from *Saccharomyces cerevisiae* S288C: a useful set of strains and plasmids for PCR-mediated gene disruption and other applications Yeast 14, 115–132

77. Winzeler, E. A., Shoemaker, D. D., Astromoff, A., Liang, H., Anderson, K., Andre, B. et al. (1999) Functional characterization of the *S. cerevisiae* genome by gene deletion and parallel analysis Science 285, 901–906

78. Wierman, M. B., Maqani, N., Strickler, E., Li, M., and Smith, J. S. (2017) Caloric restriction extends yeast chronological life span by optimizing the Snf1 (AMPK) signaling pathway. Mol Cell Biol 37, e00562–16

79. Borten, M. A., Bajikar, S. S., Sasaki, N., Clevers, H., and Janes, K. A. (2018) Automated brightfield morphometry of 3D organoid populations by OrganoSeg. Sci Rep 8, 5319

80. Ianevski, A., Giri, A. K., and Aittokallio, T. (2020) SynergyFinder 2.0: visual analytics of multi-drug combination synergies. Nucleic Acids Res 48, W488–w493

81. Yadav, B., Wennerberg, K., Aittokallio, T., and Tang, J. (2015) Searching for drug synergy in complex dose-response landscapes using an interaction potency model. Comput Struct Biotechnol J 13, 504–513

82. Shukla, S. K., Gunda, V., Abrego, J., Haridas, D., Mishra, A., Souchek, J. et al. (2015) MUC16-mediated activation of mTOR and c-Myc reprograms pancreatic cancer metabolism. Oncotarget 6, 19118–19131

83. Gunda, V., Yu, F., and Singh, P. K. (2016) Validation of metabolic alterations in microscale cell culture lysates using hydrophilic interaction liquid chromatography (HILIC)-tandem mass spectrometry-based metabolomics. PLoS One 11, e0154416

84. Moshfegh, C. M., Collins, C. W., Gunda, V., Vasanthakumar, A., Cao, J. Z., Singh, P. K. et al. (2019) Mitochondrial superoxide disrupts the metabolic and epigenetic landscape of CD4^+^ and CD8^+^ T-lymphocytes. Redox Biol 27, 101141

