## Supplemental Figures for "Chronological lifespan extension and nucleotide salvage inhibition in yeast by isonicotinamide supplementation"

**Figures S1 – S4**

**Excel files:**

**Table S1.** List of candidate genes from the INAM sensitivity screen at concentrations of 25, 50, or 75 mM.

**Table S2.** List of candidate genes from the INAM sensitivity screen at 125 mM.

**Table S3.** List of significantly enriched GO terms of the genes isolated from the 25, 50, and 75 mM INAM sensitivity screens.

**Table S5.** OASIS 2 generated statistical analysis for each CLS experiment found in the figures for this study.

**Word file:**

**Table S4.** List of yeast strains used in the study.

**Fig. S1.** Chemical genetic screen for INAM sensitive gene deletion mutants from the yeast knockout (YKO) strain collection. **A)** The haploid yeast knockout collection was pinned onto control SC plates or SC plates with increasing concentrations of INAM using a manual pinning tool. **B)** Zoomed in portion of plate 7 showing INAM sensitivity of the *tho2*Δ strain (yellow arrow). Tho2 is a transcription elongation factor. **C)** Scatter plot of scores from each deletion strain obtained in experiment 1 (x-axis) and experiment 2 (y-axis). Strains with SGAtools fitness scores lower than -0.3 in both replicates were considered sensitive to a given INAM concentration (representative graph for 75 mM INAM). **D)** Venn diagram showing the overlap of strains scored as sensitive between different INAM concentrations.

Figure S1

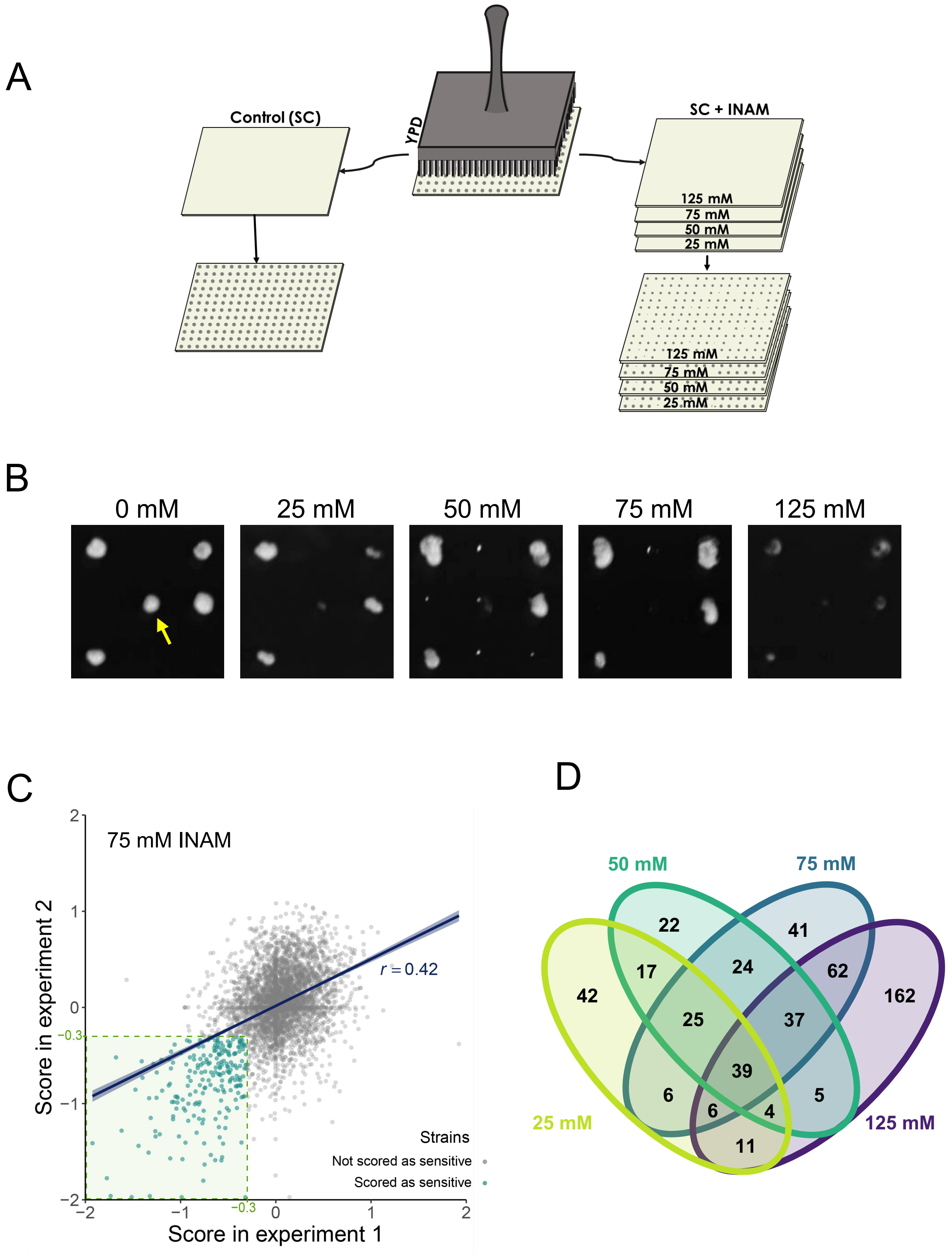
A
Control (SC)
YPD
SC + INAM
125 mM
75 mM
50 mM
25 mM
125 mM
75 mM
50 mM
25 mM
B
0 mM
25 mM
50 mM
75 mM
125 mM
C
75 mM INAM
Score in experiment 2
Score in experiment 1
r = 0.42
−0.3
−0.3
Strains
Not scored as sensitive
Scored as sensitive
D
50 mM
75 mM
25 mM
125 mM
22
41
17
24
62
42
162
25
37
39
6
6
4
5
11

**Fig. S2.** Confirmation of candidate mutants from the INAM sensitivity screen using spot test growth assays. The normalized growth scores from SGAtools are indicated for the two independent screens (performed sequentially). Scores for 25, 50, and 75 mM plates (shown) were from day 2 images, while fitness scores for 125 mM plates were from day 3 images. Fitness scores below the -0.3 cutoff are shaded in green.

Figure S2

SC | SC + 25 mM INAM | SC + 50 mM INAM | SC + 75 mM INAM

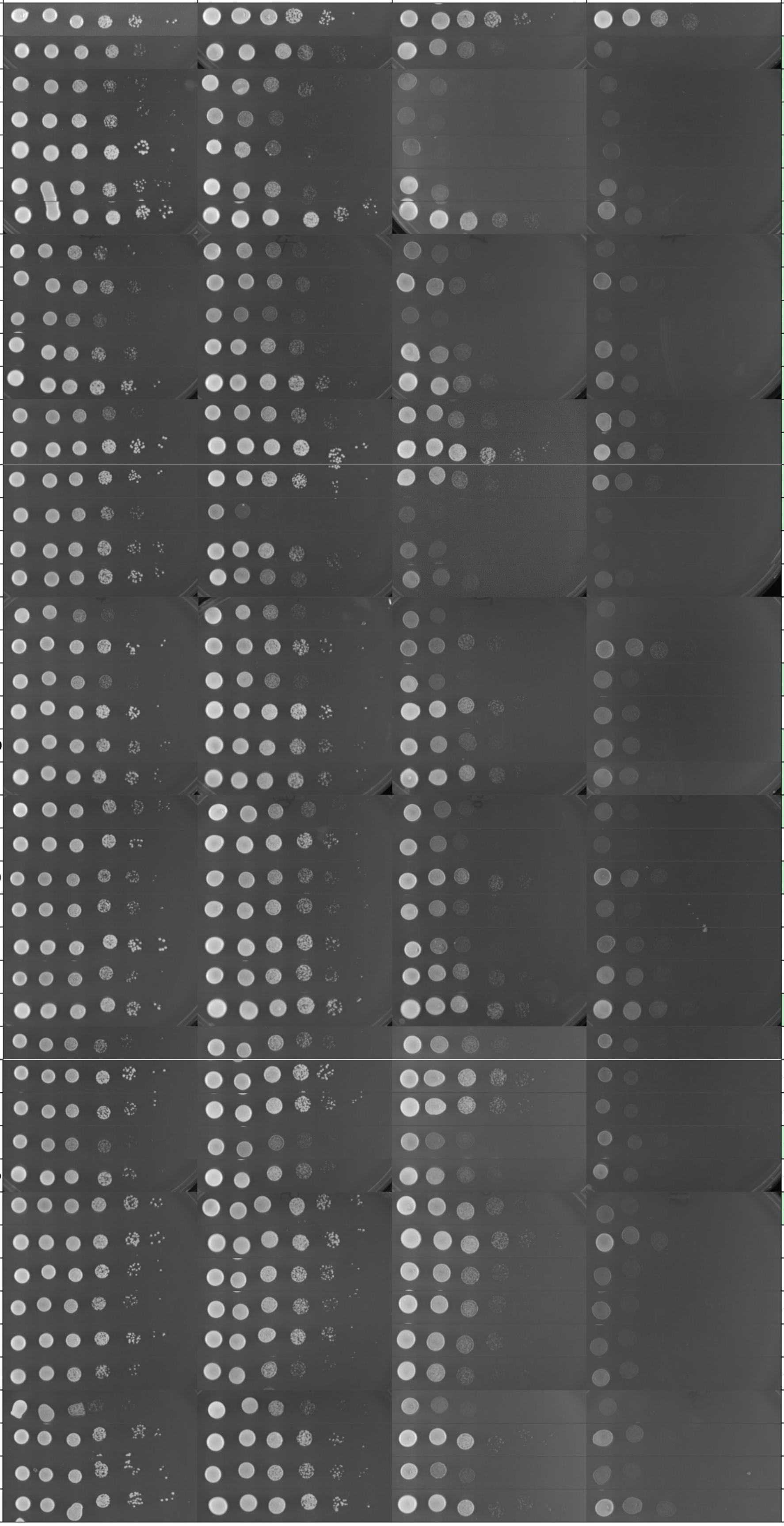

| | | Experiment #1 | | | | Experiment #2 | | | |
|---|---|---|---|---|---|---|---|---|---|
| | | Day 2 | | | Day 3 | Day 2 | | | Day 3 |
| ORFΔ | GeneΔ | 25 mM | 50 mM | 75 mM | 125 mM | 25 mM | 50 mM | 75 mM | 125 mM |
| BY4741 | | | | | | | | | |
| YOL051W | GAL11 | -0.351 | -0.443 | -0.705 | -0.705 | -0.725 | -0.725 | -0.725 | -0.758 |
| YPL129W | TAF14 | -0.345 | -0.886 | -0.845 | -0.897 | -0.669 | -0.914 | -1.144 | -1.127 |
| YNL139C | THO2 | -0.637 | -1.292 | -1.734 | -1.735 | -1.909 | -1.962 | -1.962 | -1.479 |
| YBL094C | | -1.031 | -1.031 | -1.031 | -1.031 | -0.754 | -0.968 | -0.968 | -0.939 |
| YDL117W | CYK3 | -0.445 | -0.502 | -0.696 | -0.454 | -1.349 | -1.648 | -1.648 | -0.655 |
| YDR011W | SNQ2 | -0.572 | -0.474 | -1.263 | -1.445 | -0.759 | -0.607 | -1.441 | -1.476 |
| YDR017C | KCS1 | -1.017 | -1.017 | -1.017 | -1.017 | -0.739 | -1.096 | -1.402 | -1.341 |
| YDR226W | ADK1 | -0.811 | -0.855 | -0.604 | -0.701 | -0.541 | -0.608 | -0.584 | -0.314 |
| YGL095C | VPS45 | -1.930 | -1.930 | -1.930 | -1.925 | -0.760 | -1.413 | -1.413 | -0.913 |
| YKL139W | CTK1 | -0.546 | -0.592 | -0.483 | -0.501 | -0.456 | -0.480 | -0.435 | -0.658 |
| YNL079C | TPM1 | -0.611 | -0.801 | -1.224 | -1.093 | -1.045 | -1.097 | -1.314 | -1.13 |
| YKR029C | SET3 | -0.308 | -0.322 | -0.438 | -0.512 | -0.332 | -0.332 | -0.332 | -0.344 |
| YER111C | SWI4 | -0.889 | -0.945 | -1.323 | -1.323 | -1.227 | -1.328 | -1.962 | -1.777 |
| YER139C | RTR1 | -0.592 | -0.592 | -0.592 | -0.592 | -0.939 | -1.069 | -1.069 | -0.946 |
| YLR399C | BDF1 | -0.819 | -0.819 | -0.819 | -0.807 | -0.634 | -0.634 | -0.634 | -0.468 |
| YOL012C | HTZ1 | -0.036 | -0.734 | -0.741 | -0.742 | -0.815 | -0.815 | -0.815 | -0.833 |
| YHL031C | GOS1 | -0.218 | -0.677 | -0.714 | -0.804 | -0.360 | -0.976 | -1.209 | -1.032 |
| YEL044W | IES6 | -0.125 | -1.159 | -1.384 | -1.442 | -0.061 | -0.554 | -0.742 | -0.731 |
| YER052C | HOM3 | -0.099 | -0.616 | -0.809 | -0.99 | -0.105 | -0.387 | -0.443 | -0.379 |
| YPL106C | SSE1 | -0.668 | -1.284 | -1.013 | -1.292 | -0.190 | -0.891 | -1.178 | -0.504 |
| YBR231C | SWC5 | -0.219 | -0.512 | -0.948 | -1.114 | -1.391 | -1.304 | -1.391 | -1.388 |
| YDR245W | MNN10 | -0.302 | -0.555 | -1.124 | -1.331 | 0.104 | -0.312 | -0.890 | -1.189 |
| YDR392W | SPT3 | -0.958 | -0.688 | -0.848 | -1.128 | -0.180 | -0.429 | -0.676 | -0.968 |
| YGR063C | SPT4 | -0.599 | -1.079 | -1.079 | -1.052 | -0.083 | -0.311 | -0.653 | -1.177 |
| YNL107W | YAF9 | -0.162 | -0.632 | -0.632 | -0.633 | -0.209 | -0.445 | -0.445 | -0.447 |
| YCR094W | CDC50 | -0.388 | -0.725 | -0.843 | -1.207 | -1.044 | -0.977 | -0.874 | -1.062 |
| YHL025W | SNF6 | -0.006 | -0.661 | -0.766 | -0.766 | -0.503 | -0.924 | -0.924 | -0.891 |
| YER155C | BEM2 | -0.110 | -0.752 | -0.840 | -0.95 | -1.016 | -1.016 | -1.016 | -1.009 |
| YLR337C | VRP1 | 0.023 | -1.073 | -1.441 | -1.442 | -0.052 | -1.031 | -1.443 | -1.439 |
| YLL012W | YEH1 | -0.488 | -0.001 | -1.314 | -1.905 | -0.090 | -0.447 | -1.157 | -1.535 |
| YIL128W | MET18 | -0.166 | -0.124 | -0.331 | -0.605 | -0.107 | -0.571 | -0.927 | -0.93 |
| YIL097W | FYV10 | -0.023 | 0.013 | -0.752 | -0.95 | -0.045 | -0.229 | -1.414 | -1.422 |
| YLR182W | SWI6 | -0.235 | -0.348 | -0.564 | -0.565 | -0.068 | -0.236 | -0.753 | -0.545 |
| YLR226W | BUR2 | -0.330 | -0.194 | -0.478 | -1.01 | -0.374 | -0.816 | -1.065 | -0.735 |
| YBL027W | RPL19b | -0.280 | -0.227 | -0.676 | -0.857 | -0.162 | -0.132 | -0.590 | -0.741 |
| YBR077C | SLM4 | -0.320 | -0.168 | -0.448 | -0.353 | -0.068 | -0.483 | -1.005 | -1.045 |
| YDR293C | SSD1 | 0.493 | 0.067 | -0.666 | -1.256 | -0.557 | -0.761 | -1.546 | -1.961 |
| YDR334W | SWR1 | 0.542 | 0.270 | -0.551 | -0.93 | -0.896 | -1.155 | -1.616 | -1.591 |
| YGL025C | PGD1 | -0.246 | -0.061 | -0.683 | -0.738 | 0.012 | -0.127 | -0.329 | -1.017 |
| YDR485C | SWC2 | 0.210 | -0.050 | -0.471 | -0.6 | -0.107 | -0.190 | -0.611 | -0.649 |
| YAL047C | SPC72 | -0.144 | -0.575 | -1.038 | -1.962 | 6.561 | 6.701 | -1.578 | -1.973 |
| YNL059C | ARP5 | 0.003 | -0.790 | -1.337 | -1.214 | 0.270 | -0.938 | -1.420 | -1.493 |
| YNL147W | LSM7 | 0.003 | -0.162 | -0.501 | -0.86 | -0.434 | -0.527 | -0.620 | -0.985 |
| YMR091C | NPL6 | 0.260 | -0.143 | -0.537 | -0.552 | -0.740 | -0.817 | -0.892 | -0.737 |
| YOL114C | PTH4 | 0.235 | -0.110 | -0.739 | -1.164 | -0.350 | -0.473 | -1.155 | -1.323 |

**Fig. S3.** Confirmation of additional candidate mutants from INAM sensitivity screen. **A)** Mutants that confirmed as INAM sensitive. Each experiment has its own BY4741 WT control. **B)** Mutants that did not confirm as INAM sensitive. SGAtools fitness scores for the two screens are indicated to the right, as in Fig. S2.

# A

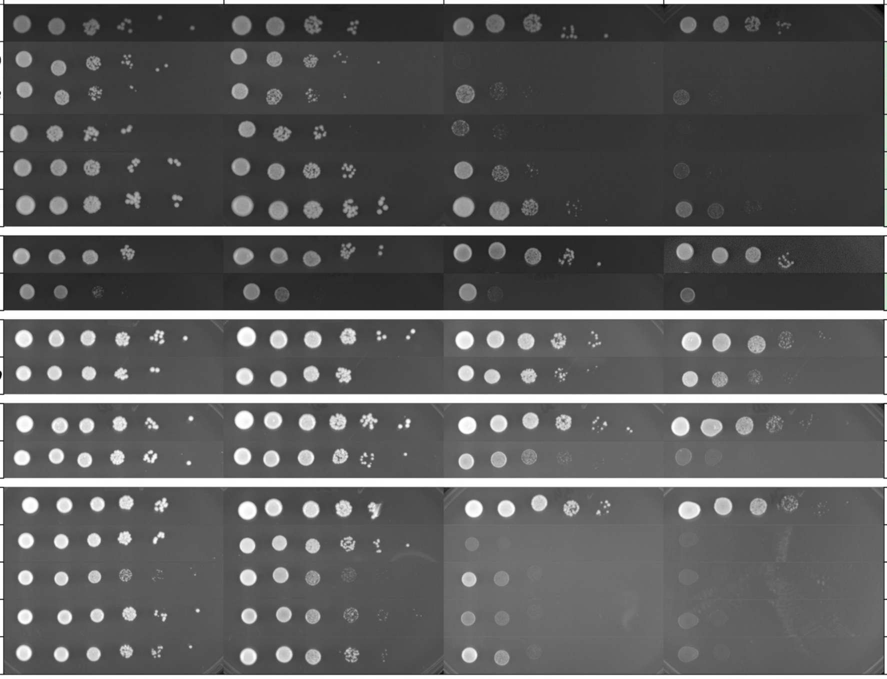


| ORFΔ | GeneΔ | Experiment #1 Day 2 25 mM | Experiment #1 Day 2 50 mM | Experiment #1 Day 2 75 mM | Experiment #1 Day 3 125 mM | Experiment #2 Day 2 25 mM | Experiment #2 Day 2 50 mM | Experiment #2 Day 2 75 mM | Experiment #2 Day 3 125 mM |
|---|---|---|---|---|---|---|---|---|---|
| BY4741 | | | | | | | | | |
| YDR364C | **CDC40** | -0.999 | -0.999 | -0.999 | -0.996 | -0.329 | -0.538 | -0.473 | -0.521 |
| *YDR532C* | ***KRE28*** | -0.481 | -0.307 | -0.654 | -0.915 | -1.454 | -1.454 | -1.417 | -0.87 |
| *YCR020W-B* | ***HTL1*** | -0.599 | -0.599 | -0.562 | -0.599 | -0.643 | -0.643 | -0.643 | -0.629 |
| *YDR138W* | ***HPR1*** | -0.555 | -0.640 | -0.897 | -0.85 | -0.029 | -0.401 | -0.774 | -0.744 |
| *YMR269W* | ***TMA23*** | -0.517 | -0.156 | -0.330 | -0.943 | -0.168 | -0.507 | -0.763 | -0.967 |
| BY4741 | | | | | | | | | |
| *YJL176C* | ***SWI3*** | -0.754 | -0.548 | -0.569 | -0.672 | -0.049 | -0.957 | -0.818 | -0.908 |
| BY4741 | | | | | | | | | |
| *YCR002C* | ***CDC10*** | 0.030 | -0.341 | -0.474 | -0.982 | -0.060 | -0.465 | -0.861 | -1.119 |
| BY4741 | | | | | | | | | |
| *YLR025W* | ***SNF7*** | 0.226 | -0.298 | -1.096 | -0.563 | -0.833 | -0.959 | -1.187 | -0.947 |
| BY4741 | | | | | | | | | |
| *YML112W* | ***CTK3*** | 0.512 | 0.191 | -0.636 | -1.112 | 0.107 | -0.493 | -1.588 | -1.244 |
| *YHL011C* | ***PRS3*** | -0.175 | -0.084 | -0.800 | -1.404 | 2.162 | -1.469 | -0.986 | -0.986 |
| *YGR155W* | ***CYS4*** | 0.865 | 0.521 | -0.320 | -1.096 | -5.046 | -5.046 | -1.973 | -1.973 |
| *YHR041C* | ***SRB2*** | -0.001 | -0.262 | -0.989 | -1.535 | 2.669 | 6.669 | -1.973 | -1.973 |

# B

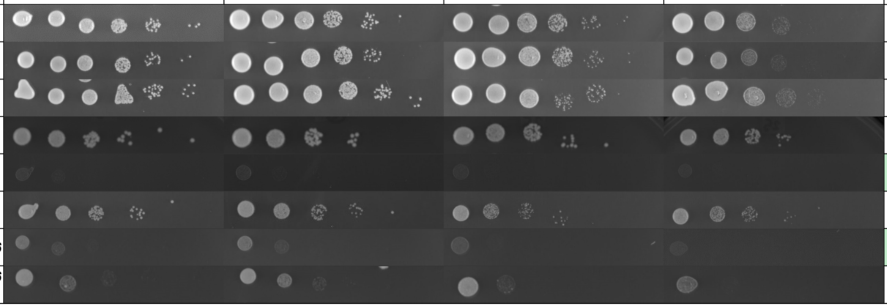


| ORFΔ | GeneΔ | Experiment #1 Day 2 25 mM | Experiment #1 Day 2 50 mM | Experiment #1 Day 2 75 mM | Experiment #1 Day 3 125 mM | Experiment #2 Day 2 25 mM | Experiment #2 Day 2 50 mM | Experiment #2 Day 2 75 mM | Experiment #2 Day 3 125 mM |
|---|---|---|---|---|---|---|---|---|---|
| BY4741 | | | | | | | | | |
| *YBL103C* | ***RTG3*** | -0.023 | 0.351 | -0.617 | -0.489 | 0.149 | 0.008 | -0.411 | -0.817 |
| *YOL115W* | ***PAP2*** | -0.159 | -0.182 | -0.325 | -0.306 | -0.392 | -0.442 | -0.401 | -0.822 |
| BY4741 | | | | | | | | | |
| *YEL046C* | ***GLY1*** | -0.926 | -1.071 | -0.605 | -0.488 | -0.852 | -0.852 | -0.852 | -0.327 |
| | ***GLY1* Day 7** | | | | | | | | |
| *YBR114W* | ***RAD16*** | -0.645 | -0.645 | -0.645 | -0.502 | -0.532 | -0.532 | -0.452 | -0.317 |
| | ***RAD16* Day 7** | | | | | | | | |

**Fig. S4.** Strains not identified by the screen but confirmed as sensitive in the spot test growth assays. These additional strains were chosen based on identity of confirmed strains from the screen and the literature.

Figure S4

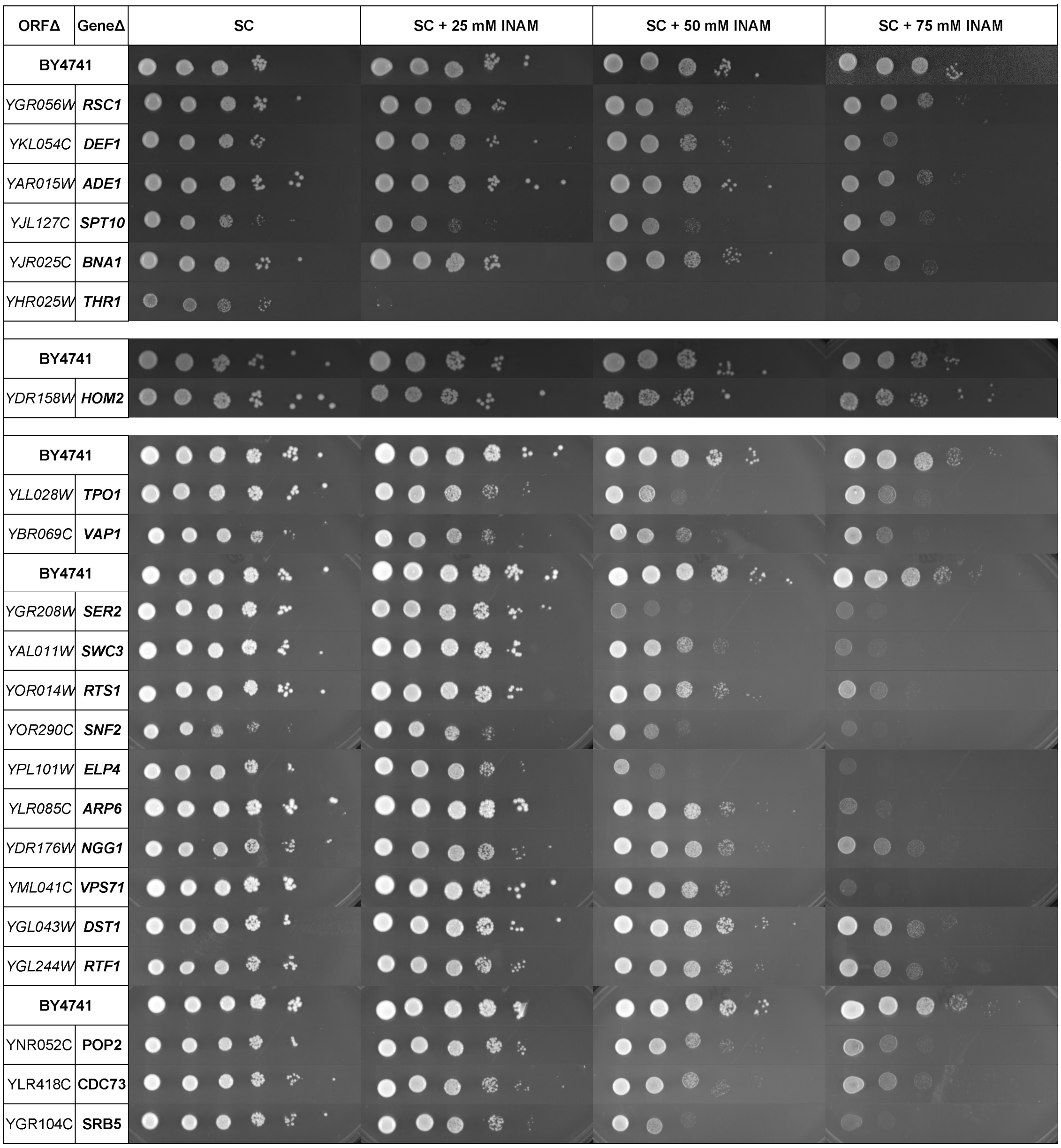
ORFΔ
GeneΔ
SC
SC + 25 mM INAM
SC + 50 mM INAM
SC + 75 mM INAM
BY4741
YGR056W RSC1
YKL054C DEF1
YAR015W ADE1
YJL127C SPT10
YJR025C BNA1
YHR025W THR1
BY4741
YDR158W HOM2
BY4741
YLL028W TPO1
YBR069C VAP1
BY4741
YGR208W SER2
YAL011W SWC3
YOR014W RTS1
YOR290C SNF2
YPL101W ELP4
YLR085C ARP6
YDR176W NGG1
YML041C VPS71
YGL043W DST1
YGL244W RTF1
BY4741
YNR052C POP2
YLR418C CDC73
YGR104C SRB5
