## Supplemental Table S4 for "Chronological lifespan extension and nucleotide salvage inhibition in yeast by isonicotinamide supplementation"

| **Table S4** | **Yeast strains used in the study** |
| --- | --- |
| **Strain** | **Genotype** |
| FY4 | *MATa* prototroph |
| BY4741 | *MATa his3Δ1 leu2Δ0 ura3Δ0 met15Δ0* |
| YPH499 | *MATa ade2-101 his3Δ200 leu2Δ1 lys2-801 ura3Δ0 trp1∆63 ura3-52* |
| SY1043 | *YPH499 made ADE2^+^* |
| YCB498 | *MATa ade2-101 his3Δ200 leu2Δ1 lys2-801 ura3Δ0 trp1∆63 ura3-52 hst1∆2::LEU2 hst2∆1::TRP1 hst3∆3::HIS3 hst4∆1::URA3 sir2∆1::URA3* |
| SY1044 | *YCB498 made ADE2^+^* |
| SY8 | *MATa his3Δ1 leu2Δ0 ura3Δ0 met15Δ0 bna1Δ::KanMX* |
| SY895 | *MATa his3Δ1 leu2Δ0 ura3Δ0 met15Δ0 bna7Δ::KanMX* |
| SY896 | *MATa his3Δ1 leu2Δ0 ura3Δ0 met15Δ0 thr1Δ::KanMX* |
| SY904 | *MATa his3Δ1 leu2Δ0 ura3Δ0 met15Δ0 gal11Δ::KanMX* |
| SY905 | *MATa his3Δ1 leu2Δ0 ura3Δ0 met15Δ0 gly1Δ::KanMX* |
| SY906 | *MATa his3Δ1 leu2Δ0 ura3Δ0 met15Δ0 taf14Δ::KanMX* |
| SY907 | *MATa his3Δ1 leu2Δ0 ura3Δ0 met15Δ0 tho2Δ::KanMX* |
| SY908 | *MATa his3Δ1 leu2Δ0 ura3Δ0 met15Δ0 rad16Δ::KanMX* |
| SY909 | *MATa his3Δ1 leu2Δ0 ura3Δ0 met15Δ0 ybl094cΔ::KanMX* |
| SY910 | *MATa his3Δ1 leu2Δ0 ura3Δ0 met15Δ0 cyk3Δ::KanMX* |
| SY911 | *MATa his3Δ1 leu2Δ0 ura3Δ0 met15Δ0 snq2Δ::KanMX* |
| SY912 | *MATa his3Δ1 leu2Δ0 ura3Δ0 met15Δ0 kcs1Δ::KanMX* |
| SY913 | *MATa his3Δ1 leu2Δ0 ura3Δ0 met15Δ0 adk1Δ::KanMX* |
| SY914 | *MATa his3Δ1 leu2Δ0 ura3Δ0 met15Δ0 cdc40Δ::KanMX* |
| SY915 | *MATa his3Δ1 leu2Δ0 ura3Δ0 met15Δ0 kre28Δ::KanMX* |
| SY916 | *MATa his3Δ1 leu2Δ0 ura3Δ0 met15Δ0 vps45Δ::KanMX* |
| SY917 | *MATa his3Δ1 leu2Δ0 ura3Δ0 met15Δ0 ctk1Δ::KanMX* |
| SY918 | *MATa his3Δ1 leu2Δ0 ura3Δ0 met15Δ0 tpm1Δ::KanMX* |
| SY919 | *MATa his3Δ1 leu2Δ0 ura3Δ0 met15Δ0 plc1Δ::KanMX* |
| SY920 | *MATa his3Δ1 leu2Δ0 ura3Δ0 met15Δ0 set3Δ::KanMX* |
| SY921 | *MATa his3Δ1 leu2Δ0 ura3Δ0 met15Δ0 htl1Δ::KanMX* |
| SY922 | *MATa his3Δ1 leu2Δ0 ura3Δ0 met15Δ0 swi4Δ::KanMX* |
| SY923 | *MATa his3Δ1 leu2Δ0 ura3Δ0 met15Δ0 rtr1Δ::KanMX* |
| SY924 | *MATa his3Δ1 leu2Δ0 ura3Δ0 met15Δ0 bdf1Δ::KanMX* |
| SY925 | *MATa his3Δ1 leu2Δ0 ura3Δ0 met15Δ0 htz1Δ::KanMX* |
| SY926 | *MATa his3Δ1 leu2Δ0 ura3Δ0 met15Δ0 gos1Δ::KanMX* |
| SY927 | *MATa his3Δ1 leu2Δ0 ura3Δ0 met15Δ0 ies6Δ::KanMX* |
| SY928 | *MATa his3Δ1 leu2Δ0 ura3Δ0 met15Δ0 hom3Δ::KanMX* |
| SY929 | *MATa his3Δ1 leu2Δ0 ura3Δ0 met15Δ0 sse1Δ::KanMX* |
| SY930 | *MATa his3Δ1 leu2Δ0 ura3Δ0 met15Δ0 swc5Δ::KanMX* |
| SY931 | *MATa his3Δ1 leu2Δ0 ura3Δ0 met15Δ0 mnn10Δ::KanMX* |
| SY932 | *MATa his3Δ1 leu2Δ0 ura3Δ0 met15Δ0 hpr1Δ::KanMX* |
| SY933 | *MATa his3Δ1 leu2Δ0 ura3Δ0 met15Δ0 spt3Δ::KanMX* |
| SY934 | *MATa his3Δ1 leu2Δ0 ura3Δ0 met15Δ0 spt4Δ::KanMX* |
| SY936 | *MATa his3Δ1 leu2Δ0 ura3Δ0 met15Δ0 yaf9Δ::KanMX* |
| SY937 | *MATa his3Δ1 leu2Δ0 ura3Δ0 met15Δ0 cdc50Δ::KanMX* |
| SY938 | *MATa his3Δ1 leu2Δ0 ura3Δ0 met15Δ0 snf6Δ::KanMX* |
| SY939 | *MATa his3Δ1 leu2Δ0 ura3Δ0 met15Δ0 bem2Δ::KanMX* |
| SY940 | *MATa his3Δ1 leu2Δ0 ura3Δ0 met15Δ0 vrp1Δ::KanMX* |
| SY941 | *MATa his3Δ1 leu2Δ0 ura3Δ0 met15Δ0 yeh1Δ::KanMX* |
| SY942 | *MATa his3Δ1 leu2Δ0 ura3Δ0 met15Δ0 tma23Δ::KanMX* |
| SY944 | *MATa his3Δ1 leu2Δ0 ura3Δ0 met15Δ0 met18Δ::KanMX* |
| SY945 | *MATa his3Δ1 leu2Δ0 ura3Δ0 met15Δ0 fyv10Δ::KanMX* |
| SY946 | *MATa his3Δ1 leu2Δ0 ura3Δ0 met15Δ0 swi6Δ::KanMX* |
| SY947 | *MATa his3Δ1 leu2Δ0 ura3Δ0 met15Δ0 bur2Δ::KanMX* |
| SY948 | *MATa his3Δ1 leu2Δ0 ura3Δ0 met15Δ0 rtg3Δ::KanMX* |
| SY949 | *MATa his3Δ1 leu2Δ0 ura3Δ0 met15Δ0 rpl19bΔ::KanMX* |
| SY950 | *MATa his3Δ1 leu2Δ0 ura3Δ0 met15Δ0 slm4Δ::KanMX* |
| SY951 | *MATa his3Δ1 leu2Δ0 ura3Δ0 met15Δ0 ssd1Δ::KanMX* |
| SY952 | *MATa his3Δ1 leu2Δ0 ura3Δ0 met15Δ0 swr1Δ::KanMX* |
| SY953 | *MATa his3Δ1 leu2Δ0 ura3Δ0 met15Δ0 pgd1Δ::KanMX* |
| SY954 | *MATa his3Δ1 leu2Δ0 ura3Δ0 met15Δ0 swc2Δ::KanMX* |
| SY955 | *MATa his3Δ1 leu2Δ0 ura3Δ0 met15Δ0 spc72Δ::KanMX* |
| SY956 | *MATa his3Δ1 leu2Δ0 ura3Δ0 met15Δ0 prs3Δ::KanMX* |
| SY957 | *MATa his3Δ1 leu2Δ0 ura3Δ0 met15Δ0 cys4Δ::KanMX* |
| SY958 | *MATa his3Δ1 leu2Δ0 ura3Δ0 met15Δ0 srb2Δ::KanMX* |
| SY959 | *MATa his3Δ1 leu2Δ0 ura3Δ0 met15Δ0 arp5Δ::KanMX* |
| SY960 | *MATa his3Δ1 leu2Δ0 ura3Δ0 met15Δ0 lsm7Δ::KanMX* |
| SY961 | *MATa his3Δ1 leu2Δ0 ura3Δ0 met15Δ0 npl6Δ::KanMX* |
| SY962 | *MATa his3Δ1 leu2Δ0 ura3Δ0 met15Δ0 pth4Δ::KanMX* |
| SY963 | *MATa his3Δ1 leu2Δ0 ura3Δ0 met15Δ0 pap2Δ::KanMX* |
| SY965 | *MATa his3Δ1 leu2Δ0 ura3Δ0 met15Δ0 ser2Δ::KanMX* |
| SY966 | *MATa his3Δ1 leu2Δ0 ura3Δ0 met15Δ0 hom2Δ::KanMX* |
| SY971 | *MATa his3Δ1 leu2Δ0 ura3Δ0 met15Δ0 srb5Δ::KanMX* |
| SY972 | *MATa his3Δ1 leu2Δ0 ura3Δ0 met15Δ0 swi3Δ::KanMX* |
| SY973 | *MATa his3Δ1 leu2Δ0 ura3Δ0 met15Δ0 rsc1Δ::KanMX* |
| SY975 | *MATa his3Δ1 leu2Δ0 ura3Δ0 met15Δ0 def1Δ::KanMX* |
| SY976 | *MATa his3Δ1 leu2Δ0 ura3Δ0 met15Δ0 ade1Δ::KanMX* |
| SY977 | *MATa his3Δ1 leu2Δ0 ura3Δ0 met15Δ0 spt10Δ::KanMX* |
| SY993 | *MATa his3Δ1 leu2Δ0 ura3Δ0 met15Δ0 tpo1Δ::KanMX* |
| SY999 | *MATa his3Δ1 leu2Δ0 ura3Δ0 met15Δ0 vap1Δ::KanMX* |
| SY1000 | *MATa his3Δ1 leu2Δ0 ura3Δ0 met15Δ0 cdc10Δ::KanMX* |
| SY1007 | *MATa his3Δ1 leu2Δ0 ura3Δ0 met15Δ0 swc3Δ::KanMX* |
| SY1008 | *MATa his3Δ1 leu2Δ0 ura3Δ0 met15Δ0 snf7Δ::KanMX* |
| SY1009 | *MATa his3Δ1 leu2Δ0 ura3Δ0 met15Δ0 rts1Δ::KanMX* |
| SY1011 | *MATa his3Δ1 leu2Δ0 ura3Δ0 met15Δ0 snf2Δ::KanMX* |
| SY1013 | *MATa his3Δ1 leu2Δ0 ura3Δ0 met15Δ0 elp4Δ::KanMX* |
| SY1014 | *MATa his3Δ1 leu2Δ0 ura3Δ0 met15Δ0 arp6Δ::KanMX* |
| SY1015 | *MATa his3Δ1 leu2Δ0 ura3Δ0 met15Δ0 ngg1Δ::KanMX* |
| SY1016 | *MATa his3Δ1 leu2Δ0 ura3Δ0 met15Δ0 dst1Δ::KanMX* |
| SY1017 | *MATa his3Δ1 leu2Δ0 ura3Δ0 met15Δ0 rtf1Δ::KanMX* |
| SY1018 | *MATa his3Δ1 leu2Δ0 ura3Δ0 met15Δ0 vps71Δ::KanMX* |
| SY1019 | *MATa his3Δ1 leu2Δ0 ura3Δ0 met15Δ0 pop2Δ::KanMX* |
| SY1020 | *MATa his3Δ1 leu2Δ0 ura3Δ0 met15Δ0 ctk3Δ::KanMX* |
| SY1124 | *MATa his3Δ1 leu2Δ0 ura3Δ0 met15Δ0 pho8Δ::KanMX* |
| SY1136 | *MATa his3Δ1 leu2Δ0 ura3Δ0 met15Δ0 pho13Δ::KanMX* |
| CL1 | *MATa his3Δ1 leu2Δ0 ura3Δ0 met15Δ0 pho13Δ::KanMX* |
